# Protein evidence of unannotated ORFs in *Drosophila* reveals unappreciated diversity in the evolution of young proteins

**DOI:** 10.1101/2022.04.04.486978

**Authors:** Eric B Zheng, Li Zhao

## Abstract

*De novo* gene origination, where a previously non-genic genomic sequence becomes genic through evolution, has been increasingly recognized as an important source of evolutionary novelty across diverse taxa. Many *de novo* genes have been proposed to be protein-coding, and in several cases have been experimentally shown to yield protein products. However, the systematic study of *de novo* proteins has been hampered by doubts regarding the translation of their transcripts without the experimental observation of protein products. Using a systematic, ORF-focused mass-spectrometry-first computational approach, we identify almost 1000 unannotated open reading frames with evidence of translation (utORFs) in the model organism *Drosophila melanogaster*, 371 of which have canonical start codons. To quantify the comparative genomic similarity of these utORFs across *Drosophila* and to infer phylostratigraphic age, we further develop a synteny-based protein similarity approach. Combining these results with reference datasets on tissue- and life-stage-specific transcription and conservation, we identify different properties amongst these utORFs. Contrary to expectations, the fastest-evolving utORFs are not the youngest evolutionarily. We observed more utORFs in the brain than in the testis. Most of the identified utORFs may be of *de novo* origin, even accounting for the possibility of false-negative similarity detection. Finally, sequence divergence after an inferred *de novo* origin event remains substantial, raising the possibility that *de novo* proteins turn over frequently. Our results suggest that there is substantial unappreciated diversity in *de novo* protein evolution: many more may exist than have been previously appreciated; there may be divergent evolutionary trajectories; and *de novo* proteins may be gained and lost frequently. All in all, there may not exist a single characteristic model of *de novo* protein evolution, but rather complex origins and evolutionary trajectories for *de novo* proteins.

**Impact statement:** The analysis of mass-spectrometry data for all possible open reading frames reveals protein evidence for evolutionarily young, unannotated proteins with distinct characters.

## Introduction

While the study of the evolution of entire species or the fixation of individual mutations has matured over decades, our understanding of how genes evolve is relatively less mature. The dominant mechanism is duplication and divergence (Ohno, 1970; Long et al., 2003), but the early years of the genomic era revealed a seeming paradox of species-specific “orphan genes”. These genes lacked homologs even in closely related species, which made it necessary to invoke non-standard duplication and divergence mechanisms (Domazet-Loso and Tautz, 2003). Advances in both the sequencing of genomes and in comparative genomics soon led to an explosion of work on the origins of orphan genes, generally identified through lineage-specificity.

Lineage-specific orphan genes can originate from a variety of mechanisms, ranging from extreme, rapid divergence after duplication (Chen et al., 2010; Domazet-Loso and Tautz, 2003) to horizontal gene transfer (Verster et al., 2019) to *de novo* gene birth. In the latter mechanism, previously non-genic genomic sequence acquires genic function, thus yielding a *de novo* gene that is characterized by its lineage-specificity (Begun et al., 2006; Levine et al., 2006; McLysaght and Hurst, 2016; Schlötterer, 2015; Van Oss and Carvunis, 2019). *De novo* genes are identified from diverse taxa from fruit flies (Begun et al., 2006; Levine et al., 2006), to yeast (Cai et al., 2008) and even humans (Knowles and McLysaght, 2009; Xie et al., 2012). While it would seem unlikely for such *de novo* genes to gain viable function from presumably unoptimized sequence, empirical and population genetics studies have demonstrated that they have important functions (Cai et al., 2008; Lange et al., 2021; Levine et al., 2006; Reinhardt et al., 2013; Xie et al., 2019; Zhao et al., 2014). However, exactly what proportion of lineage-specific genes are contributed through *de novo* gene birth remains actively studied (Tautz and Domazet-Lošo, 2011; McLysaght and Guerzoni, 2015; Vakirlis et al., 2020).

Despite significant interest, the study of evolutionarily young genes is surprisingly conceptually difficult. While an empirical investigation of biological effect is ideal, designing such experiments for a specific evolutionarily young gene can be challenging due to the likely lack of homology to previously studied genes. Even identifying a given candidate genetic locus as an evolutionarily young gene can be fraught with challenges: evolutionary youth is functionally equivalent to concluding lineage-specificity, but methods like phylostratigraphy via BLAST (Domazet-Loso et al., 2007), while useful and intuitive, can be vulnerable to false-negative detections, especially for short genes (Moyers and Zhang, 2015, 2016).

Moreover, while researchers generally agree that a gene should be functional, exactly how this should be defined and assessed can vary substantially (Van Oss and Carvunis, 2019). As an example, whether evolutionarily young genes function as RNAs or as proteins is often an area of debate. One of the earliest recognized young genes is an RNA (Wang et al., 2002), other studies also provided such support (Cridland et al., 2022; Ruiz-Orera et al., 2015); in contrast, in several cases even *de novo* genes have been empirically shown to yield protein products (Bungard et al., 2017; Cai et al., 2008; Knowles and McLysaght, 2009; Li et al., 2010; Zhang et al., 2019). While defining a gene as one that can code for a translated protein is convenient and intuitive, evidence for the translation into polypeptides of lineage-specific genes and in particular, *de novo* genes as a class, is often lacking (Van Oss and Carvunis, 2019). As a result, much effort remains needed for systematic approaches to studying evolutionarily young genes and proteins.

Proteomics approaches like shotgun mass spectrometry (MS) can be a powerful tool for the rapid identification of translational evidence for many potential proteins at once, and would be a good fit for such a systematic study. However, standard shotgun MS analysis approaches require *a priori* assumptions about the sequences of the proteins to be detected (Sinitcyn, Rudolph, and Cox 2018). In the case of discovering possible lineage-specific proteins, this *a priori* requirement can be confounding, as lineage-specific proteins may be systematically under-annotated due to their lack of homology.

Here, we employ a systematic, ORF-focused MS-first computational approach, to identify unannotated open reading frames with evidence of translation (utORFs) in *Drosophila melanogaster*. We further develop a synteny-based protein similarity approach to quantify the comparative genomic similarity of these utORFs across *Drosophila*, to estimate their phylostratigraphic age, and finally to infer possible *de novo* origin for the vast majority of these utORFs. Combined with reference datasets on tissue- and life stage-specific transcription and conservation, this data suggests differences amongst these utORFs with respect to their evolutionary and functional properties. Finally, we independently support a subset of these utORFs via additional MS empirical data and ribosome profiling data.

## Results

### A comprehensive database of all possible open reading frames in the *Drosophila melanogaster* genome

As a first step towards the systematic discovery of unannotated proteins in *Drosophila melanogaster*, we generated a comprehensive database of all possible open reading frames (ORFs) in the *D. melanogaster* genome. This comprehensive database is the result of translating the entire annotated genome in all six frames; we also included the annotated transcriptome to enable discovery of non-canonical translation phenomena such as alternate frame ORFs (de Klerk and ’t Hoen 2015). In this database, each potential ORF is defined as a contiguous span of amino acid-coding codons; so, these potential ORFs do not necessarily begin with or contain a canonical start codon. This is necessary in order to account for unannotated splicing events, as a six-frame translation of the genome cannot correctly associate the exons of a gene. Since no coding exon within the coding sequence of a gene can contain a stop codon, the set of all codon spans between stop codons is a superset of the set of all exons in the genome. In addition to annotated ORFs (FlyBase r6.15 release), this comprehensive potential ORF database contains 4,582,998 unique potential ORFs that are not otherwise annotated. Of these additional potential ORFs, the median length is 32 amino acids and the 95th percentile is 96 amino acids. Considering solely the 2,597,794 potential ORFs that contain a canonical start codon, the median length following the start codon remains 32 amino acids, and the 95th percentile is 112 amino acids. In total, this suggests that the *D. melanogaster* genome has substantial latent potential for generating new protein-coding genes. Of course, the overwhelming majority of these potential ORFs are neither transcribed nor translated and are thus not functional (Durand et al. 2019), but considering these will allow us to scan the potential proteomic diversity in *D. melanogaster*.

### Two-round MS search improves discovery of unannotated ORFs without compromising FDR

A comprehensive proteome of *D. melanogaster* was recently published (Casas-Vila et al. 2017), including data from 15 stages of the whole lifecycle of the fruit fly (ranging from 0-2h embryos to adults, including larval and pupal stages), as well as sharper time-resolution data covering 14 stages of embryogenesis (hourly timepoints from 0-6 hours; bi-hourly timepoints from 6 to 20 hours). We analyzed the *Casas- Villa et al.* data by searching for the complete potential ORF database described above to discover evidence of translation of unannotated ORFs in *D. melanogaster*. Because the number of potential ORFs is several orders of magnitude greater than the number of annotated proteins, the search space is enormously expanded, and thus a traditional one-round search is likely to miss a very large number of identifications at a given false-discovery rate (FDR). Accordingly, to improve total sensitivity while maintaining an acceptable FDR, we used two rounds of analysis (**Methods**). We identify dramatically more novel unannotated protein groups with statistical significance in this two-round approach than in a traditional single-round search, while the number of false positive identifications is comparable (**Supplemental File 1A**). The substantial enrichment (around ten-fold) of unannotated ORFs relative to either the observed number of false positives or the desired FDR strongly supports the validity of the two-round approach.

In total, after additional filtering (see **Methods**), our two-round search identified 3123 unannotated unique peptides that underlie 993 unannotated ORFs without homology to annotated proteins (**Figure 1A; Figure 1B; Figure 1–Source Data 1**; **Figure 1–Source Data 2**), which we hereafter refer to as unannotated translated ORFs (utORFs). utORFs range in length from 14 amino acids (aa) to over 100 aa, with the longest being 1748 aa (**Figure 1C**). The median length of a utORF is 37 aa, and the mean is 48.6 aa. Among these utORFs, 331 proteins have canonical start codons with 1027 peptides that originate from sequence after the canonical start codon, implying that on average each protein has 3 supporting peptide-spectrum matches. The mean and median lengths of a utORF starting with a canonical start codon are 53.5 aa and 37 aa, respectively, which is very similar to utORFs overall. Unless specifically mentioned, we analyze henceforth the set of 993 utORFs and not the 331 with canonical start codons.

**Figure 1.**
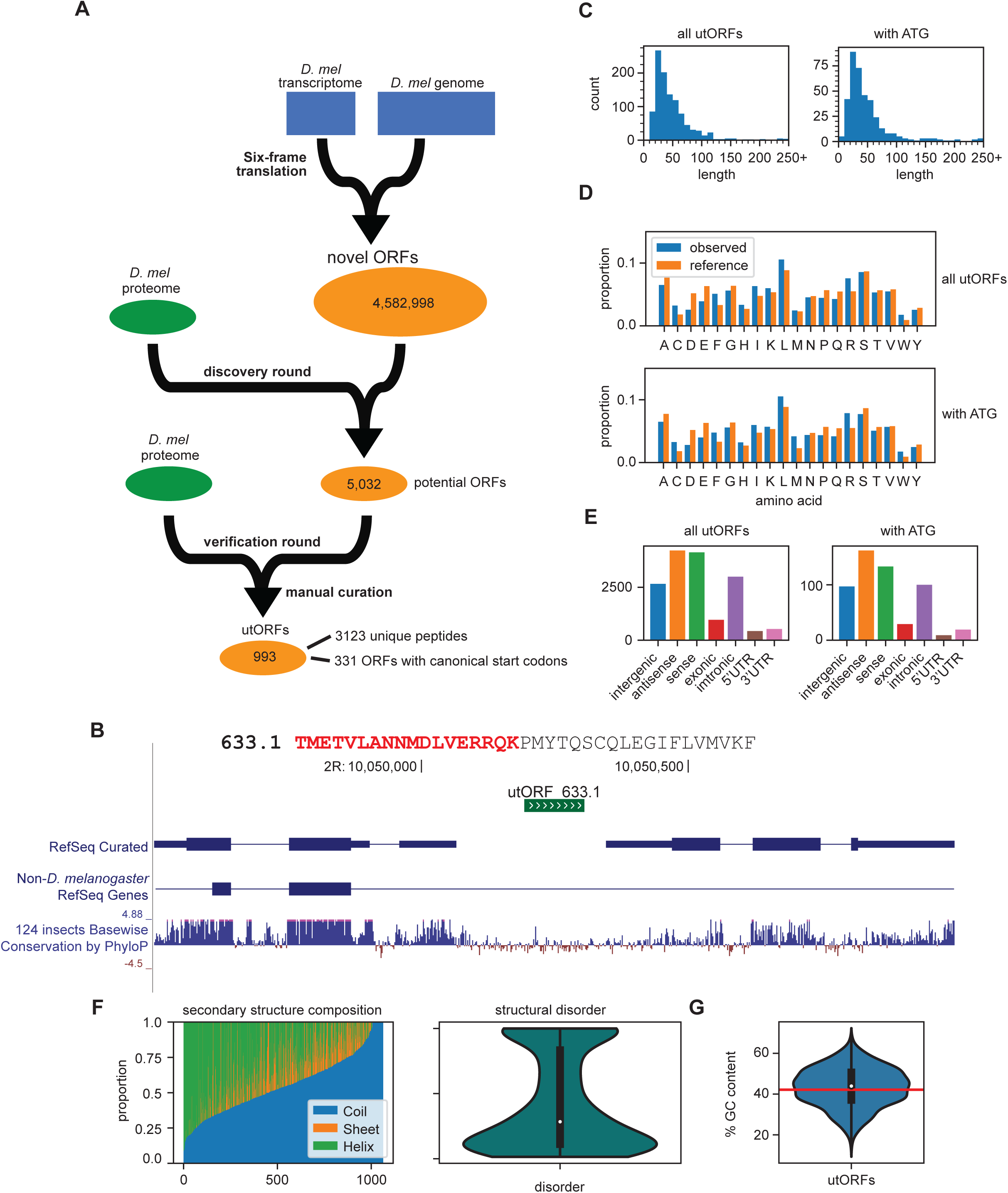
Identification of utORFs and their properties. **A**. We detected utORFs by searching via a two-round approach through a comprehensive database of potential ORFs that was generated from a six-frame translation of the transcriptome and genome (see Methods). **B**: an example utORF. The full sequence of utORF 633.1 is shown. The peptide supporting the inference of this utORF is bolded and colored red. **C**: distribution of lengths of utORFs. **D**: distribution of amino acids in utORFs (blue) and in the reference D. melanogaster proteome (orange). Standard single-letter abbreviations for amino acids are used. **E**: genomic locations of utORFs. Since a utORF can overlap multiple items, e.g., an ORF that overlaps with an annotated intron, exon, and another gene antisense to the first, categories are not exclusive. **F**: left: secondary structure composition of utORFs. Approximately half of utORFs have a coiled-coil secondary structure, with the remainder a mixture of beta sheets and alpha helices. right: predicted structural disorder of utORFs. **G**: Percent GC content of utORFs and GC content of the X and autosomes of the *D. melanogaster* genome (red line).

Compared to the annotated *D. melanogaster* proteome, utORF sequences have similar amino acid composition (**Figure 1D**; Spearman rank-order correlation of amino acid frequencies, utORFs to annotated proteome: 0.767, *p* = 7.98 E -5). The amino acid composition for those utORFs with start codons is also highly similar. Some hydrophobic and positively-charged amino acids, such as leucine, arginine, and tryptophan are overrepresented relative to annotated proteins, while negatively-charged amino acids, such as aspartate and glutamate are underrepresented. Overall, the distribution of positively-charged to negatively-charged amino acids is significantly different among utORFs versus annotated *D. melanogaster* protein sequences (Chi-squared test with Yates’s correction, Chi-sq = 1463.55, df=1, *p* < machine epsilon). In addition, the distribution of hydrophobic amino acids versus polar amino acids is also highly significantly different (same test, Chi-sq = 44.96, df=1, *p* < 1E -10). These differences may be explained by the amino acid composition of potential ORFs, as the implied amino acid composition of all potential ORFs is more similar to utORFs than annotated proteins (Spearman rank-order correlation of amino acid frequencies, potential ORFs to utORFs: 0.970, *p* = 1.71 E -12; potential ORFs to annotated proteome: 0.785, *p* = 4.15 E -5).

All 993 utORFs have unique genomic locations (**Figure 1–Source Data 3**). Unannotated translated ORFs reside in a range of genomic locations, including intergenic, intronic, or UTRs (**Figure 1E**). About one-third of utORFs are located intergenically; of these, about one-third again are longer than 50 aa and contain a canonical start codon. We propose that these may be single-exon unannotated proteins. A considerable number of utORFs overlap with annotated genes, with an even distribution between sense and anti-sense directions relative to the annotation. Of the former, several are located either upstream or downstream of the canonical CDS; these may be instances of uORFs or translation read-through, which have been previously described as having important biological functions (Andrews and Rothnagel 2014). Notably, with the exception of a modest reduction in the relative proportion of sense utORFs, the distribution of locations is not different for those utORFs with a canonical start codon. The genomic sequences that make up utORFs on average have greater GC content compared that of the overall *D. melanogaster* genome (X and autosomes; mean utORF GC content 43.4% vs. X and autosome GC content 42.15%; 1-sample two-tailed *t*-test, *p*<1E-4), but they have wide variety, with some with as much as 62% GC content and others as few as 21% (**Figure 1G**). Intriguingly, the GC content of annotated genes (annotated gene GC content 42.9%) is between that of the genome and utORFs, and utORFs do not have significantly different GC content compared to annotated genes (1-sample two-tailed *t*-test, *p* > 0.05). This suggests either that a selective pressure may act towards increasing the GC content in utORFs, or that utORFs may be more likely to originate from regions with higher GC content.

Approximately half of utORFs (566 of 993; 57.0%) have a majority coiled-coil secondary structure, with the remainder a mixture of beta sheets and alpha helices. Notably, this remains true even when normalizing for length, as the weighted proportion of sequence in a predicted coiled-coil structure is on average about half (**Figure 1F**). Predictions of structural disorder of utORFs suggest that while they are rather disordered, most retain a substantial proportion that is ordered. The median proportion of disordered utORFs is 24.5%.

### Novel peptides identified are not artifacts caused by polymorphism in conserved genes

It is possible that polymorphisms within a population can yield unique peptides due to amino acid substitutions, thus resulting in inaccurate attribution of such peptides to utORFs (Faridi, Purcell, and Croft 2018). However, such polymorphisms can contribute at most a small fraction of the peptides that support the identified utORFs. To quantify this, using a trie, we calculated the minimum distance between the peptides that uniquely support utORFs and the expected set of peptides from the reference proteome (see **Methods**). Of the 3123 tryptic peptides that are uniquely identified from utORFs, only 1 could possibly result from a single amino acid substitution of a tryptic peptide from an annotated protein; none could result from a single amino acid insertion or deletion. A simultaneous double amino acid substitution could explain an additional 5 peptides; 24 more peptides could result from a double insertion or deletion. Thus, single or double amino acid substitutions can explain at most 30 of 3123 unique tryptic peptides (< 1%). As a result, it is exceedingly likely that the peptides detected here are the product of utORFs rather than polymorphisms at annotated genes.

### Inferring ages of unannotated translated ORFs

In order to compare the conservation of utORFs across species, it is necessary to find the orthologous sequence with near base-pair-level resolution. Due to the algorithmic compromises necessary for whole-genome searches, BLAST homology searches do not guarantee that they will return optimum results, especially for short sequences that may be highly diverged. Moreover, any homology search that solely relies on searching for a given query sequence necessarily neglects broader genomic context like synteny. Conversely, common genomic utilities like UCSC’s *liftOver* are explicitly not recommended for fine-detailed genomic coordinate conversions. To address these issues, we used a pre-existing multiple-sequence-alignment (MSA) to guide a maximally-optimal local genome alignment (**Figure 2 AB**). This allows the combination of both whole genomic contexts as well as the algorithmic guarantees of the Smith-Waterman alignment.

**Figure 2.**
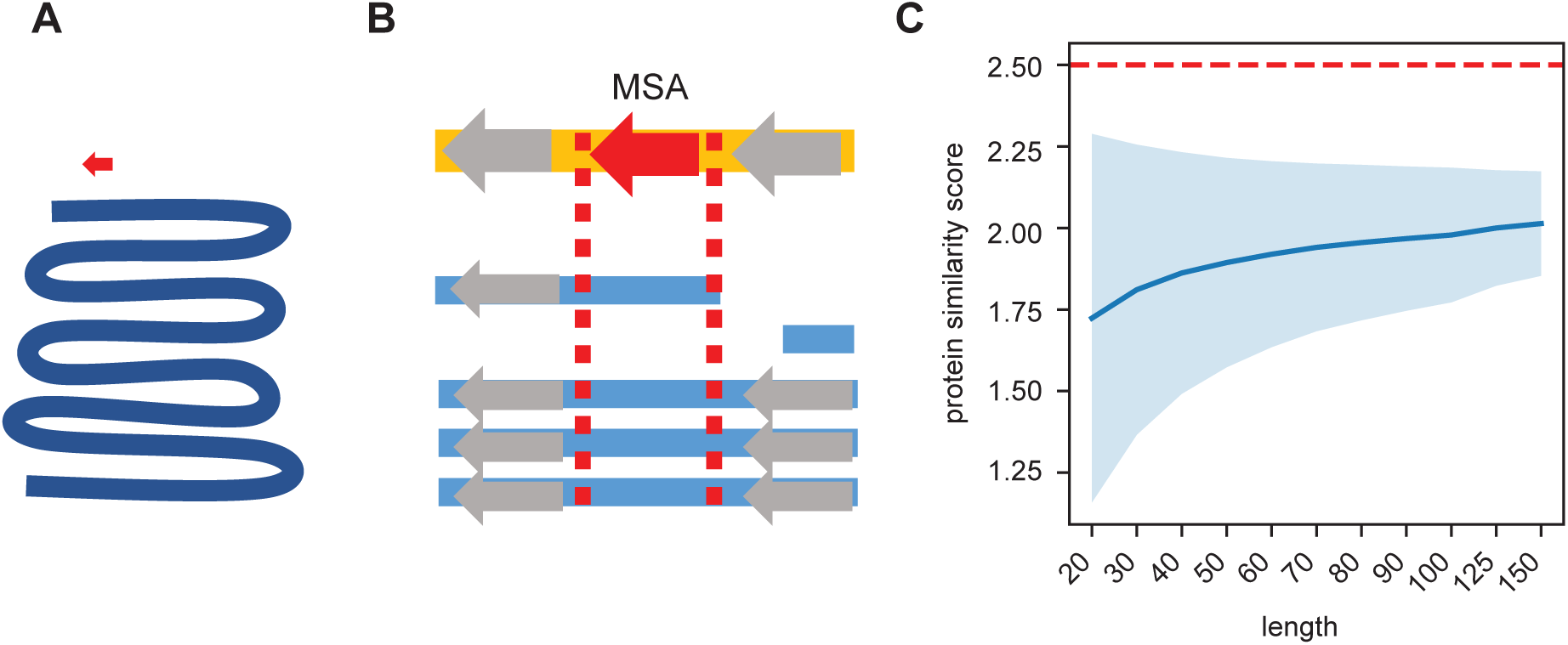
Synteny-based orthology detection and protein sequence similarity quantification. **A:** When performing a simple homology search for a locus of interest (red arrow) across a given genome (blue line), the search space is orders of magnitude larger, requiring heuristic shortcuts to reduce the computational complexity. **B:** Using a multiple sequence alignment, we can simply find the block that contains the locus of interest and use that to evaluate potential orthologs in other genomes. The search space is approximately a similar size as that of the original locus, so an optimal search is computationally feasible. Syntenic information (i.e., flanking genes, gray arrows) is encoded within the multiple sequence alignment. **C:** We calculated the pairwise protein similarity score (see Methods) across 10000 pairs of randomly-generated sequences of lengths 20-150 with amino acid frequencies matching that of the annotated *D. melanogaster* proteome. The mean score (blue line) and 2 standard deviations (shading) are shown. Our significance threshold of 2.5 lies beyond 2 standard deviations from expectation.

To estimate a null model of protein similarity scores, we calculated control distributions with varying assumptions. Assuming amino acids drawn from the distribution of amino acids in the annotated *D. melanogaster* proteome, 10000 simulated pairs of randomly-generated protein sequences have a mean expected similarity score ranging from 1.7 to 2.0 (**Figure 2C**). We chose a conservative threshold of 2.5 points as a threshold for determining significant sequence similarity, as it is approximately at least 2 standard deviations above the mean expectation.

We used the above-described protein similarity score to identify significant orthologs across related *Drosophila* and insect species using the UCSC 27-way MSA (**Figure 3A**). We inferred gene ages for each utORF, where the inferred origin of a utORF corresponded to the last common ancestor between *Drosophila melanogaster* and the most distantly related species where a significant ortholog was identified. We find that utORFs have a wide range of inferred gene ages, ranging from utORFs specific just to *D. melanogaster* to utORFs that may be conserved broadly through ancient insect taxa (**Figure 3B**). The plurality of utORFs likely predate the last common ancestor of *Drosophila* with the most-distant significant ortholog in *D. grimshawi* or any of its sister species; the next largest group predate the last common ancestor of the *Drosophila* species group, with the most-distant significant ortholog in *D. ficusphila* or its sister species. Notably, gene age inferences are not dramatically affected by choice of significance threshold (**Figure 3–figure supplement 1**).

**Figure 3:**
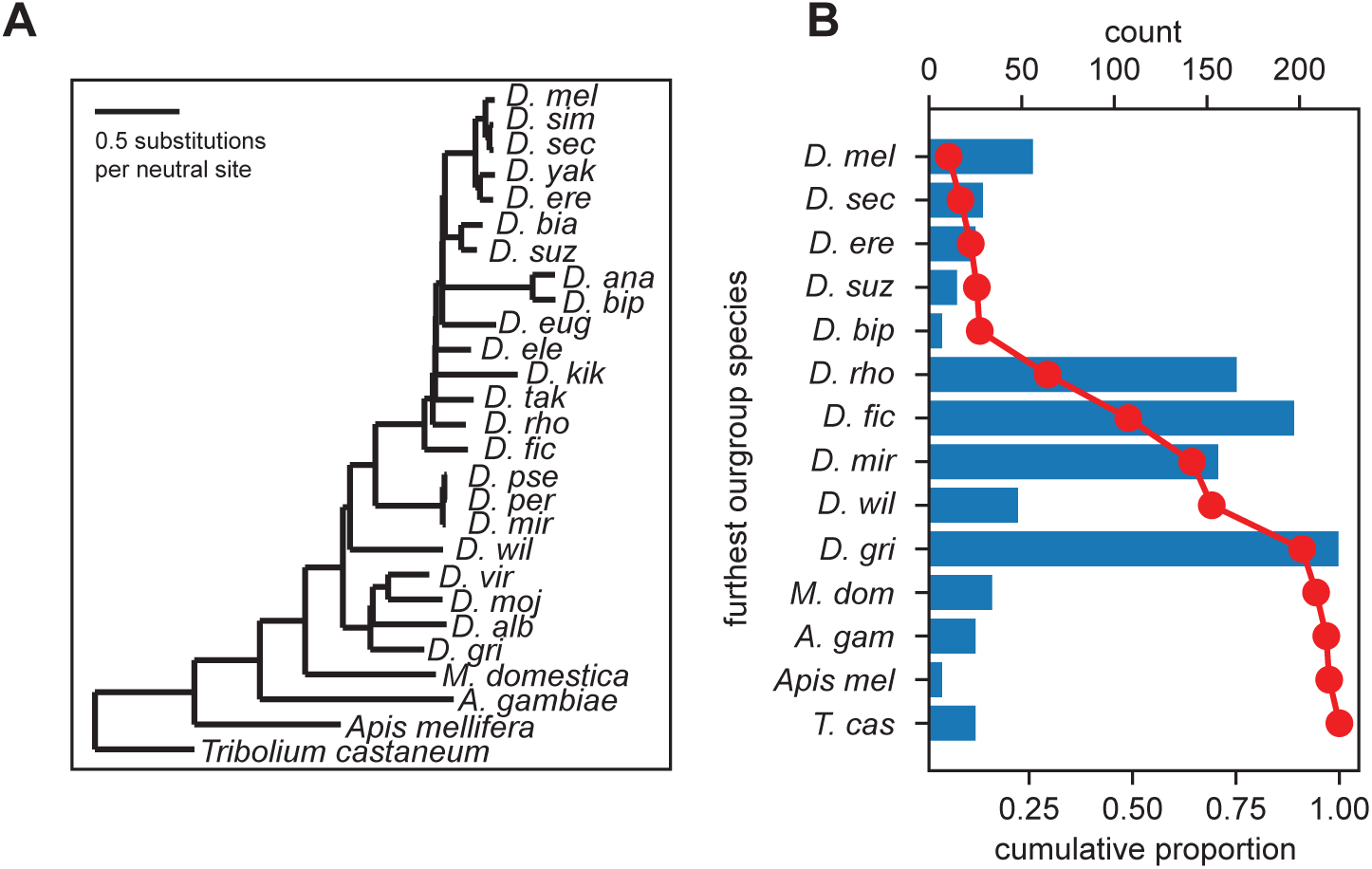
Inferred gene ages of the utORFs. **A**: The reference phylogenetic tree used for these analyses (UCSC 27-way insect alignment). Abbreviations are as follows: *D. mel*: *Drosophila melanogaster*, *D. sim*: *D. simulans*, *D. sec*: *D. sechellia*, *D. yak*: *D. yakuba D. ere*: *D. erecta*, *D. bia*: *D. biarmipes*, *D. suz*: *D. suzukii*, *D. ana*: *D. ananassae*, *D. bip*: *D. bipectinata*, *D. eug*: *D. eugracilis*, *D. ele*: *D. elegans*, *D. kik*: *D. kikkawai*, *D. tak*: *D. takahashii*, *D. rho*: *D. rhopaloa*, *D. fic*: *D. ficusphila*, *D. pse*: *D. pseudoobscura*, *D. per*: *D. persimilis*, *D. mir*: *D. miranda*, *D. wil*: *D. willistoni*, *D. vir: D. virilis*, *D. moj: D. mojavensis*, *D. alb: D. albomicans*, *D. gri*: *D. grimshawi*, *M. dom*: *Musca domestica*, *A. gam*: *Anopheles gambiae*, *Apis mel*: *Apis mellifera*, *T. cas*: *Tribolium castaneum*. **B**: The most distantly related species in which a significant ortholog of a utORF exists varies. The red line illustrates the cumulative distribution of loci. For convenience, sister species are grouped together under one species (e.g., *D. virilis* with *D. grimshawi*; *D. eugracilis*, *D. elegans*, *D. kikkawai*, and *D. takahashii* with *D. rhopaloa*; etc.). Abbreviations as in **A**.

### Latent class analysis reveals subpopulations of unannotated translated ORFs

Latent class analysis (LCA) is a statistical method that supposes that the population under study is comprised of a mixture of distinct subpopulations – the “latent” classes (Collins and Lanza 2009). Importantly, this is the only major assumption necessary, so the analysis is robust to the shape of distributions or correlations between variables. Moreover, LCA can be used both deductively and inductively; its primary output is a table of probabilities for categorical variables conditional on class membership. Thus, these conditional probabilities can be used deductively to describe differences in the inferred classes, and they can be used inductively to predict class membership given an individual set of observations.

Due to the unbiased strategy of identification, it is likely that the unannotated translated ORFs identified here consist of a mixture of loci resulting from different, distinct evolutionary processes, ranging from intergenic *de novo* genes to alternative open reading frames of conserved annotated genes (Samandi et al. 2017). In addition, certain key variables, like inferred gene age and genomic location, are inherently categorical. Rather than simple inter-variable correlations, LCA is a valuable lens for understanding the unannotated loci identified here.

Accordingly, latent class analysis suggests that the population of high-confidence unannotated translated ORFs can be subdivided into five classes (Methods; **Table 1; Figure 4-Source Data 1**) and interpretations assigned based off differences in the various factor probabilities conditional on class membership (Collins and Lanza 2009).

**Table 1:** Latent class analysis of all unannotated translated ORFs.

| Class | Interpretation | Estimated Percent | Number |
| --- | --- | --- | --- |
| 1 | putatively nonfunctional loci | 4.35% | 41 |
| 2 | melanogaster-specific ORFs | 5.71% | 54 |
| 3 | fast-evolving ORFs | 12.03% | 96 |
| 4 | unannotated ORFs | 57.61% | 591 |
| 5 | alternative frame ORFs | 20.30% | 161 |

Class 1 loci are notable for minimal transcription (**Figure 4C**) as well as a marked bias of being intergenic or antisense (**Figure 4A**). They also are almost exclusively of intermediate length between 20 and 50 residues (**Figure 4B**). This supports an interpretation as putatively nonfunctional loci. Class 2 loci are remarkable for monophyly (**Figure 4C**), likely due to very recent inferred emergence (**Figure 4D**), suggesting that these may be *melanogaster*-specific loci. Dramatic differences in overall conservation (**Figure 4E**) as well as subtler ones in inferred emergence times (**Figure 4D**) distinguish classes 3, 4, and 5. Class 5 is also remarkable for strong bias towards locations overlapping existing, annotated genes, as well as high conservation, supporting an interpretation as ORFs in alternative reading frames. Finally, class 3 can be distinguished from class 4 through a more recent inferred age of emergence approximately at the last common ancestor of the *melanogaster* subgroup (**Figure 4D**) and through remarkably low conservation that is even more frequent than that of *melanogaster*-specific ORFs. This supports a distinction between class 3 as fast-evolving ORFs and class 4 as general unannotated ORFs.

**Figure 4.**
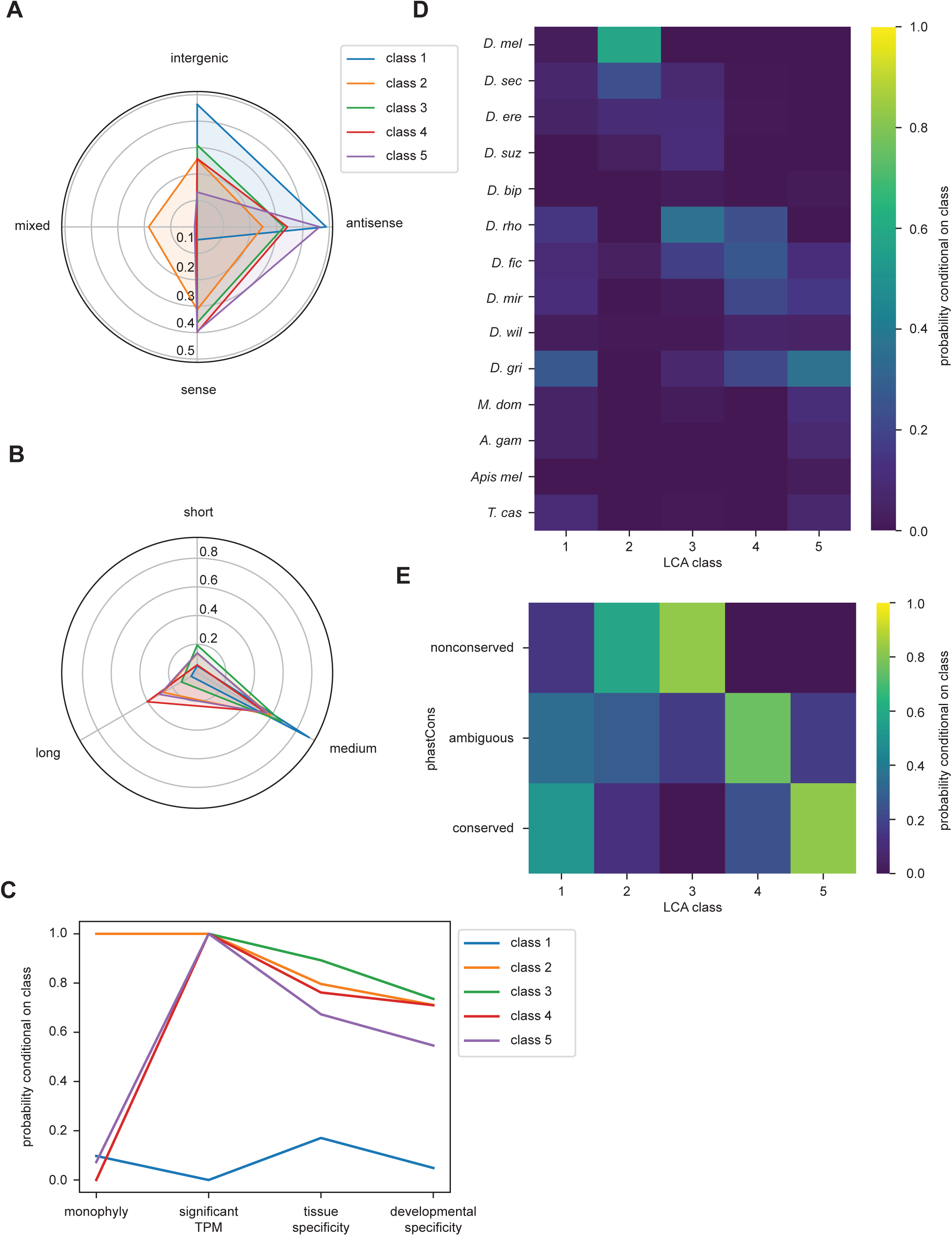
Latent class analysis of the utORFs reveals differences between classes. **A**: Class 1 is notably distinct for strong bias towards intergenic and antisense locations at the expense of sense locations. Class 2 is notable for being relatively unbiased, and for being the only class with appreciable members in a combination of locations. Class 5 is strongly skewed towards antisense and sense locations. **B**: Class 1 is almost exclusively of intermediate length. Class 4 has the greatest length bias, followed by class 2. tiny: fewer than 20 residues; medium: from 20 to 49 residues; long: 50 or more residues. **C**: Class 1 is notably distinct from the others for minimal transcription. Low tissue and developmental specificity may be an effect of minimal transcription. Class 2 is remarkable for being entirely monophyletic. Class 5 has slightly lower tissue and developmental specificities than classes 2-4. sigTPM: maximum per-sample TPM 0.1; tau: tissue specificity 0.8; dev_tau: developmental specificity 0.8. **D**: Class 2 is by far the youngest. Class 3 tends to be of an intermediate age, with inferred emergence at around the LCA of the *melanogaster* subgroup (droRho). In contrast, class 4 emergence is distributed throughout the LCA of the *melanogaster* subgroup and the *Drosophila* genus. **E**: Class 2 is notable for overall low conservation. Class 3 is remarkably even less conserved. Class 4 and 5 are distinguished through differences in intermediate vs. significant conservation. unconserved: phastCons score < 0.2; amb: phastCons score ≥ 0.2 and < 0.8; significant: phastCons score ≥ 0.8.

With inferred classes in hand, it is possible to use the classes for forward inferences by assigning each utORF to a class and comparing the classes. Importantly, while binning continuous variables is necessary for conducting latent class analysis, comparisons between classes can be done on the original continuous variables, removing that potential source of bias, and additional variables not originally used for assigning classes can be used as well. The posterior probabilities of class membership for each utORF suggest generally clean partitioning between classes, as each utORF generally has a high posterior probability of belonging to only a single class (**Figure 4–figure supplement 1**). The exceptions are utORFs with membership in either class 4 (unannotated ORFs) or class 5 (alternative-frame ORFs). However, allowing an additional sixth class leads to overfitting (see **Methods**). Accordingly, we use the maximum posterior probability per utORF to assign class membership for further analysis.

For certain parameters, differences by inferred class membership recapitulate expected trends. For example, phastCons conservation scores vary by class, with the lowest conservation in fast-evolving and *melanogaster*-specific utORFs (**Figure 5A**). Alternative-frame utORFs are very well conserved due to the additional constraints imposed by their overlapping annotated genes. Despite not being included as a variable in LCA, phyloP conservation scores also vary significantly by class (**Figure 5B**). Negative phyloP scores are consistent with faster evolution; 61.4% (27 of 44) are from fast-evolving utORFs and 24.1% (15 of 44) are from *melanogaster*-specific utORFs. Importantly. while phastCons and phyloP scores are significantly correlated, only phyloP scores can differentiate between relative low conservation and accelerated evolution. This recapitulation of faster evolution supports the overall latent class analysis approach. Finally, consistent with the assigned interpretation, *melanogaster*-specific and fast-evolving utORFs are the youngest according to phylostratigraphy (**Figure 5C**). Thus, the fastest-evolving utORFs, which are the fast-evolving utORFs with negative phyloP scores, are surprisingly not the youngest utORFs, as the distinct *melanogaster*-specific utORFs are younger.

**Figure 5.**
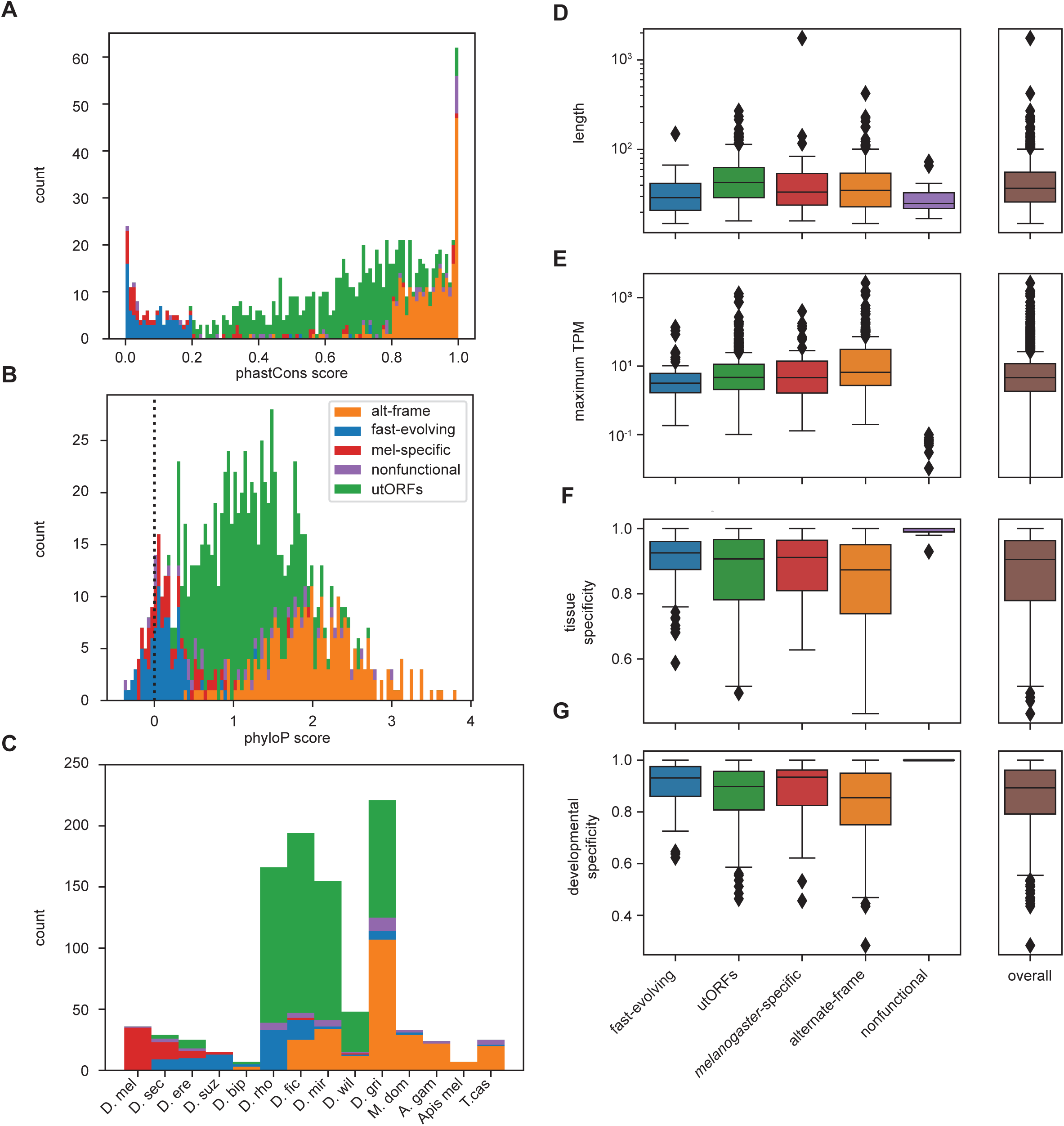
Differences between inferred classes recapitulate expected trends in age and conservation and reveal surprising trends in lengths and expression. **A**: As expected, phastCons conservation scores vary by class. Scores near 0 indicate low conservation while scores near 1 indicate high conservation. Note that fast-evolving and mel-specific loci have dramatically lower conservation whereas alt-frame loci are very well conserved. **B**: Despite not being included as a variable in LCA, phyloP conservation scores also vary by class. Scores below 0 indicate potential fast evolution (acceleration) while scores above 0 indicate conservation. Note that the lowest scores predominantly occur in fast-evolving and mel-specific loci, and that alt-frame loci remain the best-conserved. **C**: Distribution of phylostratigraphically-inferred gene ages by inferred latent class. **D**: Class variations in length are generally minimal, with the exception of utORFs being slightly longer than average, and fast-evolving and nonfunctional loci being shorter than average. **E**: Maximum observed transcription across all FlyAtlas2 and modENCODE samples varies by class. **F**: Tissue specificity (tau) calculated from FlyAtlas2 data shows that nonfunctional and fast- evolving loci are the most specific, with alternate frame loci the least. **G**: Developmental specificity (tau) calculated from modENCODE data similarly shows that nonfunctional and fast-evolving loci are the most specific, with alternate frame loci the least.

For other parameters, differences between inferred classes reveals surprising trends. Fast-evolving utORFs and putatively nonfunctional utORFs are significantly shorter than all others (one-sided Brunner-Munzel nonparametric test of lengths vs. all other utORFs, *p* << 0.001 for both). Interestingly, *melanogaster*-specific utORFs are not significantly shorter (same test, *p*=0.11), despite prior reports that younger genes tend to be shorter. With regards to maximum observed transcription, alternative-frame utORFs tend to be significantly better-transcribed than all other utORFs (one-sided Brunner-Munzel nonparametric test of log-transformed TPMs vs. all other utORFs, *p* << 0.001). This is likely due to active transcription of the overlapping pre-existing loci. In contrast, fast-evolving utORFs are significantly less well-transcribed (same test, *p* < 0.001), while differences in *melanogaster*-specific and utORFs are statistically insignificant. Consistent with these trends, alternative-frame utORFs are less tissue- and developmentally-specific (one-sided Brunner-Munzel test of tau across FlyAtlas2 tissues, *p* < 0.001; same test of tau across modENCODE developmental stages, *p* < 0.001) while fast-evolving utORFs are more specifically-expressed (tau across FlyAtlas2 tissues, *p* = 0.0039; tau across modENCODE developmental stages, *p* < 0.001). Unusually, a plurality of fast-evolving utORFs (33 of 96, 34.4%) have their highest expression in adult fly tissues in the brain, with the next most common location the testis (10 of 96, 10.4%). Moreover, while transcription of annotated genes is high in both the brain and testis, fast-evolving utORFs are dramatically less-transcribed in the testis than in the brain (75th percentile log_10_ of TPM in brain, fast-evolving utORFs: 0.172; vs. annotated genes: 1.320; in testis, fast-evolving utORFs: - 0.662; vs. annotated genes 1.177; delta log_10_ of TPM in testis versus in brain, fast-evolving utORFs vs. annotated genes: one-sided Mann-Whitney U test *p* = 7.5E-6; **Figure 5–figure supplement 1**). Indeed, with the exception of alternate-frame utORFs, most utORFs are not transcribed in the testis but are transcribed in the brain, while most annotated genes are highly transcribed in both (**Figure 5–figure supplement 1**). This is somewhat intriguing as previous work suggests that evolutionarily young genes – particularly *de novo* genes – emerge in the testis.

When considering only utORFs with canonical start codons, the results are generally qualitatively similar (**Figure 4–figure supplement 2, Figure 5–figure supplement 2**). Five latent classes remain the best fit, as allowing a sixth class has unfavorable AIC and BIC effects. Class assignments can proceed with similar logic; however, utORFs with start codons are longer overall (**Figure 4–figure supplement 2B**), so some class interpretations change somewhat. Class 3 appears biased towards intergenic locations and longer length, so we assign it an interpretation of “intergenic ORFs” (**Supplemental File 1B**).

Partitioning between classes remains good (**Figure 4–figure supplement 3**). Turning once again towards forward inferences, we recapitulate the patterns seen for the conservation of *melanogaster*-specific and alternative-frame ORFs. While differences in length are apparent between classes (**Figure 5–figure supplement 2D**), they are not statistically significant; the *melanogaster*-specific class still is not significantly shorter (one-sided Brunner-Munzel nonparametric test of lengths vs. all other utORFs: intergenic ORFs, longer, *p* = 0.11; alternate-frame, shorter, *p* = 0.15; *melanogaster*-specific, shorter, *p* = 0.36; putatively nonfunctional, shorter, *p* = 0.20). Finally, utORFs with canonical start sites remain less transcribed in the testis than the brain (delta log base 10 of TPM in testis versus in brain, utORFs with canonical start sites vs. annotated genes: one-sided Mann-Whitney U test *p* = 7.3E-21). Non-canonical start initiation is increasingly recognized, especially for short ORFs; evidence for translation initiation at sites other than AUG has come from both ribosome profiling and mass spectrometry approaches from a variety of model species (Chen et al., 2020; Ingolia et al., 2011; Ma et al., 2014; Wu et al., 2020). Since we recapitulated similar results for utORFs with canonical start sites as for all utORFs, we continued with all utORFs.

### Many unannotated translated ORFs have evidence consistent with a *de novo* origin

A genetic locus emerging through *de novo* gene birth from nongenic sequence would be expected to be detected in a lineage-specific manner with detectable genomic homology despite evidence inconsistent with functional conservation. That is, a utORF may be of *de novo* origin if there exists at least one species where an orthologous ORF is not significant despite genomic homology. Ideally, the strongest inferences of *de novo* origin would rely on multiple outgroup species. This protects against an erroneous *de novo* inference due to a homology detection failure rather than true functional changes, which has been extensively described as a potential source of erroneous identifications (Moyers and Zhang 2015, 2016, 2018). Many of our utORFs have such stringent evidence consistent with a *de novo* origin (**Figure 6A**), with at least two supporting outgroups originating from different phyla. Between 40 and 75% of identified loci across all inferred classes may be *de novo*, with the greatest proportion in the fast-evolving, *melanogaster*-specific, and unannotated ORFs. These differences between classes are statistically significant (Pearson’s Chi-squared test, 4 degrees of freedom, *X*^2^ = 31.941, *p* = 1.97E-6). For all utORFs with potential *de novo* origin, 394 (67.8%) have 3 or more such supporting outgroup species. The number of such supporting outgroup species appears to be more affected by the availability of such than by genomic divergence, as the youngest classes (*melanogaster*-specific and fast-evolving) have more supporting outgroup species than the comparatively older alternative-frame class (**Figure 6B**).

**Figure 6.**
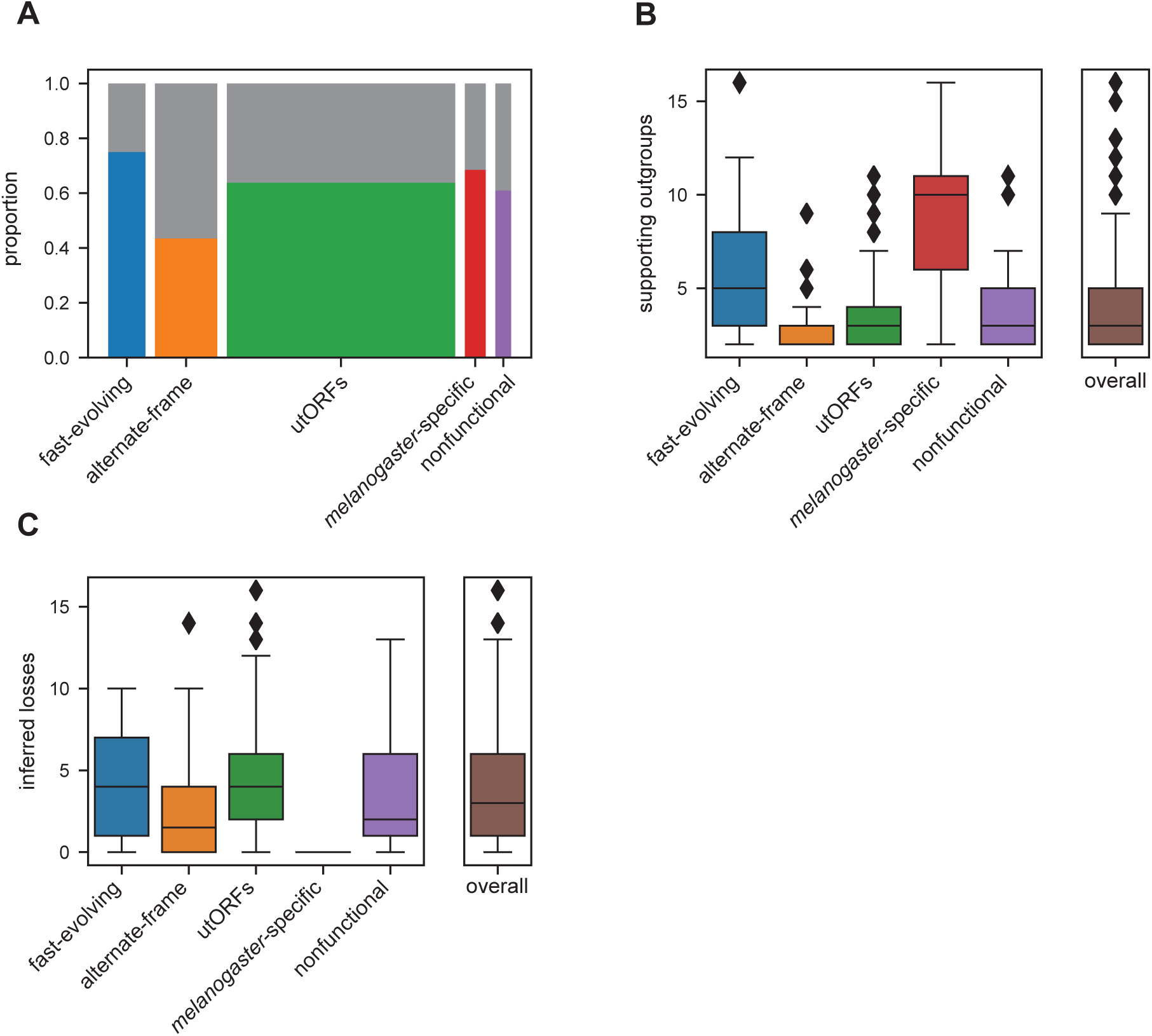
Many utORFs have evidence consistent with a de novo origin. **A**: Proportion of utORFs by inferred class with genomic conservation consistent with *de novo* origin. Box widths correlate with size of class (**Table 1**). **B**: Number of supporting outgroups by inferred class for loci with potential *de novo* origin. **C**: Number of losses inferred after potential de novo origin.

Finally, we considered the frequency by which utORFs of potential *de novo* origin is lost. Since we consider the protein sequence to be equivalent to the function, we use similar logic as for finding support for *de novo* origin, but looking only at species descended from the inferred potential de novo origin event. We find that losses of utORFs of potential *de novo* origin are extremely common; for all utORFs with potential *de novo* origin, 470 (80.9%) have at least 1 potential inferred loss (**Figure 6C**).

### Observed retention times of many peptides underlying unannotated translated ORFs are similar to those of peptides underlying annotated proteins

One possible weakness of the methodology used in this study is that peptide-spectrum matches (PSMs) are prone to false identifications (Nesvizhskii and Aebersold 2005). One method that has been proposed to evaluate PSMs is to use the retention time of identified peptides on the liquid chromatography system upstream of the mass spectrometer; retention time should vary as a function of the peptide’s sequence. While calculating retention time of a given peptide *a priori* remains an unsolved problem, substantial progress has been made on predicting retention times of sets of peptides *a posteriori*. To evaluate the validity of PSMs of utORFs, we used AutoRT (Wen *et al*. 2020) to predict retention times of peptides supporting annotated FlyBase proteins as well as utORFs. While some peptides supporting utORFs have drastically larger differences from the predicted retention times (**Figure 7–figure supplement 1**), a sizeable number have observed retention times that are similar to peptides from annotated FlyBase proteins. Specifically, 385 of 3123 peptides (12.3%) supporting utORFs have at least one PSM with a delta retention time that is smaller than the 95th percentile of the delta retention times of all PSMs that support annotated FlyBase proteins; within this subset, the difference between predicted and observed retention times is modest (**Figure 7A**). These peptides are sufficient to support 270 utORFs, where melanogaster-specific ORFs are slightly more likely and nonfunctional ORFs somewhat less likely to be supported (**Supplemental File 1C**); however, this observation is not statistically significant (Chi-squared test with 4 degrees of freedom; Chi-sq = 4.45; p > 0.05).

**Figure 7.**
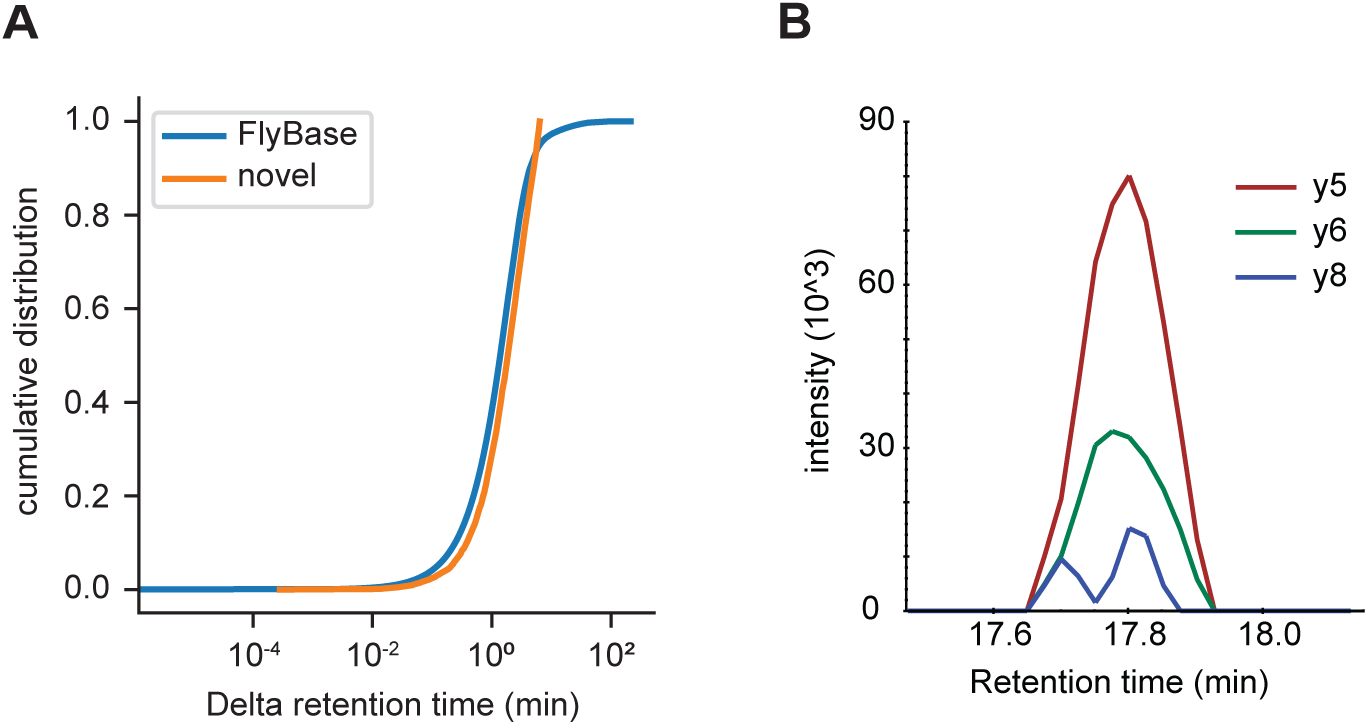
Independent validation of utORF identification. **A:** Cumulative distribution of differences between observed and predicted retention times for peptide-spectrum matches of peptides supporting annotated FlyBase proteins (orange) and peptide-spectrum matches of peptides supporting utORFs that are no worse than the 95th percentile of annotated FlyBase PSMs (blue). **B:** Transitions detected via PRM of the peptide GPGAAISQR from protein extracts of *D. melanogaster* embryos.

### utORFs can be independently validated

To further validate that the discovered utORFs are not artifactual, we collected our own data to develop independent empirical support for these utORFs. We collected targeted MS data from mixed 0-24 hour *D. melanogaster* embryos (RAL517 strain), targeting a manually-curated set of 15 target utORFs chosen both for suitability for targeted MS (e.g. few missed cleavages; no post-translational modifications) as well as expression in embryo from modENCODE data. Of these 15 targeted utORFs, we were able to detect supporting evidence for 1 through the observation of 3 transitions (**Figure 7B**). We also collected shotgun MS data from mixed 0-24 hour embryos; we were able to detect supporting evidence for 3 additional utORFs from the shotgun MS data at a FDR of 1% (**Figure 7-Source Data 1**); this shotgun dataset was fairly shallow, as we identified 1963 annotated proteins compared to the approximately 8000 protein groups reported by Casas-Vila et al. (2017). Given the low sensitivity of MS, having evidence for utORFs from independent datasets supports an unneglectable translation level for at least a subset of the utORFs. Furthermore, we re-analyzed published ribosome profiling data (Patraquim et al. 2020; Zhang et al. 2018) and identified supporting evidence from an additional 243 utORFs. Of these, alternate-frame ORFs are unsurprisingly the most frequently supported, perhaps due to translation of the canonical ORF, but the other classes are well-represented with the exception of the putatively nonfunctional utORFs (**Figure 7–figure supplement 2**). The biases across classes are statistically significant (Pearson’s Chi-squared test, 4 degrees of freedom, *X*^2 = 41.569, *p* << 0.001), and they are consistent with our proposed interpretation of the classes. Finally, we specifically looked for evidence of framing in ribosome profiling data by implementing the binomial statistical test used for the first report of framing in *Drosophila* (Patraquim et al., 2020). Of 81 utORFs with more than 2 unique footprints, 17 utORFs (21.0%) have statistically-significant biases towards the observed frame in at least 1 stage of embryogenesis, supporting the notion of active translation (alpha 0.05; 11 utORFs at alpha 0.001).

Together, this evidence supports our identification of the pool of utORFs overall. We accordingly integrated all data that we analyzed with respect to additional lines of evidence supporting our utORFs and factors of interest in order to identify utORFs of highest priority for downstream functional analyses; we used a combination of high transcription from RNAseq atlases, supporting validation from either our MS experiments or framed ribosome profiling reads, and conservation metrics. By calculating ranks across each of these factors and averaging ranks per utORF, we generated a priority list for our utORFs (**Figure 7 - Source Data 2**). Of the top 50 utORFs, 24 are alternate-frame ORFs, 14 unannotated ORFs, 7 melanogaster-specific ORFs, and 4 fast-evolving ORFs. This is similar to the overall proportion of class assignments, except alternate-frame ORFs are somewhat over-represented due a higher incidence of framed ribosome profiling reads. These utORFs may be the most promising candidates for future experimental work.

## Discussion

### Identification of hundreds of potential protein products of *de novo* origin

In this work, we identify almost one thousand possible utORFs from a systematic, ORF-focused MS-first computational approach. These utORFs are unlikely to be annotation failures due to the lack of sequence homology to annotated proteins. Instead, like many similar “orphan genes”, these may be of *de novo* origin: the vast majority have a syntenic sequence conservation pattern that is consistent with *de novo* origin (**Figure 6AB**). The total number of identifications is larger than previous studies of *de novo* genes within a relatively short timescale in *Drosophila* (Begun et al. 2006; Chen, Zhang, and Long 2010; Zhao et al. 2014), and is consistent with the identification of thousands of possible *de novo* polypeptides from systematic studies in other biological systems (Ruiz-Orera et al. 2018; Stein et al. 2018; Durand et al. 2019). Moreover, while evidence of translation of proposed *de novo* genes is generally lacking (McLysaght and Hurst 2016), in this case, the method of identification allows a stronger inference of translation.

Calling *de novo* origin for a given gene is challenging, as the call is almost wholly dependent on negative results in the outgroup lineages. For example, in small phylogenies with a single outgroup, it is possible for a single false negative call to lead to an incorrect inference of *de novo* origin. Recent studies have demonstrated that false negatives may be alarmingly likely, especially when using simple BLAST-based phylostratigraphy (Moyers and Zhang 2015, 2016). Here, our method of calling *de novo* origin employs proposed best practices of incorporating syntenic information (Arendsee et al. 2019; Vakirlis and McLysaght 2019) and is robust to individual sequence homology calls, improving our confidence in our inferences. Importantly, our inferences of gene age and potential *de novo* origin do not rely at all on the absence of sequence from the MSA, which would be analogous to the lack of a significant BLAST hit.

Instead, our inferences of *de novo* origin rely on protein similarity scores below our threshold of significance (**Figure 6B**), which ensures that these inferences are not simply due to extensive genomic divergence beyond the limit of detection. If anything, our formulation of the protein similarity score may be vulnerable to calling spurious similarity due to shared genomic ancestry between closely related species. However, such errors would cause us to infer systematically older *de novo* origin dates for each gene; given that we still conclude that many utORFs are of potential *de novo* origin, the impact of such possible errors is low.

Other mechanisms beyond *de novo* origin could potentially produce “orphan” / lineage-specific genes. For example, horizontal gene transfer from microorganisms like viruses has been reported in *Drosophila* (Verster et al., 2019), and much attention has been focused on sequence divergence (e.g. after pseudogenization) as an alternative explanation for lineage-specific genes (Moyers and Zhang, 2015 Weisman et al., 2020). When, as is often the case for our utORFs, there are many supporting outgroups, multiple pseudogenization events are necessary by parsimony to produce the observed pattern; while this seems unlikely, we cannot exclude the possibility of repeated substantial divergence. However, work explicitly focusing on the possibility that sequence divergence is responsible for lineage-specific genes recently concluded that extensive divergence was not likely the dominant mechanism (Vakirlis et al., 2020).

Our confidence in our identifications of utORFs is supported by our confirmatory results from mass spectrometry and published ribosome profiling, including the observation of statistically significant framing from ribosome-profiling data. While the absolute number of recapitulated utORF detections is relatively low, mass spectrometry has notoriously low power, even in well-studied proteomes like humans (Baker et al. 2017). Thus, it is not particularly surprising that we were unable to recapitulate most utORFs in a relatively heterogenous sample, as we expect utORFs to be of low abundance and potentially only expressed under specific time- and tissue- limited conditions. Unlike sequencing approaches, the empirical readout of mass spectrometry is not directly mappable to information (i.e. the underlying nucleotide sequence), so false identifications of peptide-spectrum matches are a notorious problem in the field (Nesvizhskii and Aebersold, 2005). However, the peptides underlying our identification of utORFs are not artifacts caused by polymorphism in conserved genes. Moreover, our results suggest that the reverse-decoy approach for controlling the number of false-positive identifications remains valid for our two-round method: the power for the identification of annotated Drosophila genes is nearly the same between a standard MS approach and our two-round approach (**Supplemental File 1A**). Further functional investigation of utORF products would be a potential extension of this work (**Figure 7 - Source Data 2**).

### Class inference reveals heterogenous evolutionary dynamics

In contrast to previous work, our approach of applying latent class analysis allows additional refinement beyond that achievable by simple per-variable analyses. For example, previous studies have described an association between gene age and length: namely, that younger genes tend to be shorter (Carvunis et al. 2012; Neme and Tautz 2013; Wilson et al. 2017). Here, we find that the youngest class of utORFs (the *melanogaster* specific class) are not significantly shorter; instead, the fast-evolving and the nonfunctional classes are the shortest overall (**Figure 5C**). Importantly, while length is an input into the latent class analysis, it is not the most obvious differentiator between the classes, nor is it the primary basis upon which we interpreted the classes. Since the fast-evolving class is still quite young, it is possible that previous reports of an association between gene age and length are related to a conflation of fast evolution with gene age. Importantly, our analysis is not able to evaluate whether there is a direction in the historical evolutionary trajectory of the length of a given utORF. Thus, we cannot distinguish between the possibilities that shorter utORFs are faster evolving or that faster evolving ORFs require a shorter length. This question merits further examination in future.

Previous studies, including our own, have suggested that evolutionarily young genes tend to be expressed in the testis (Levine et al. 2006; Begun et al. 2007; Soumillon et al. 2013; Zhao et al. 2014; Luis Villanueva-Cañas et al. 2017; Witt et al. 2019), though other tissues, like the brain, can give rise to evolutionarily young genes (Chen et al., 2012). In contrast to that, the utORFs identified here are expressed more in the brain than in the testis (**Figure 5–figure supplement 1**) in each inferred class.

Moreover, while most annotated genes are highly expressed in the testis, most utORFs are not expressed at all in the testis, yet they are expressed in the brain (**Figure 5–figure supplement 1**). Considering that most previously-reported *de novo* genes also tend to be expressed in the testis (Levine et al. 2006; Begun et al. 2007; Zhao et al. 2014; Luis Villanueva-Cañas et al. 2017; Witt et al. 2019), our results may suggest that the proportion of protein-coding *de novo* genes might be higher in brain than in testis, or that our MS- first approach allows the detection of otherwise-missed protein-coding *de novo* genes. Furthermore, these utORFs may have biologically significant effects in the brain. For example, the maximum observed expression of a plurality of the relatively short, fast-evolving ORFs is in the brain, suggesting that some could act as neuropeptides.

Despite the success of our latent class analysis approach in elucidating distinct evolutionary trajectories, half of our identified utORFs of interest remain consolidated in a single class. However, additional classes cannot be safely supported due to overfitting. Additional approaches may be necessary in order to further characterize these utORFs.

### Turnover in utORFs may be frequent

While *de novo* gene birth has been increasingly well-studied over recent years, *de novo* gene death has been examined less-closely. It is obvious that *de novo* gene death must be relatively frequent, as otherwise the net size of proteomes would be unrealistically large relative to the annotation. Consistent with this intuition, we observe that most *de novo* genes diverge by enough to be indistinguishable from random sequences in at least part of the clade descended from the last common ancestor after the *de novo* gene birth event (**Figure 6C**). Since we consider conservation of the protein-coding sequence to be approximately equivalent to the function of the locus, we consider such sequence divergence as evidence of gene death.

An implication of most models of *de novo* gene origins is that *de novo* gene birth is rare at the individual gene level and thus occurs only once in evolutionary history. However, the very large number of gene death events suggests a second possibility. It is possible that the rates of gene birth and death may be more balanced; that is, instead of being born once and recurrently lost, a subset of *de novo* genes may be born, die, and subsequently reborn (Schmitz et al., 2018; Stein et al., 2018; Durand et al., 2019). Thus, some *de novo* gene evolution may not proceed unidirectionally, but rather like a random walk along a continuum. This is consistent with the proto-gene model (Carvunis et al. 2012) or the pre-adaptation model (Wilson et al. 2017). Under the proto-gene model, the frequent deaths of our utORFs would imply that they are proto-genes and not genes. However, with the exception of the *melanogaster*-specific utORFs, our utORFs are often conserved across species with significant divergence times, which implies that some may have evolutionary constraints and may in fact be functional genes. It is difficult to make strong conclusions here without getting trapped in the philosophical quagmire of the nature of “geneness”; nevertheless, it is possible that no single “grand unified theory” of *de novo* gene origination can describe all phenomena.

### Outlook

Together, our results show that evolution of young proteins may progress along different, distinct trajectories in *Drosophila*. Whether similarly distinct trajectories are apparent in other model species such as yeast or mammals remains to be seen. Of note, *Drosophila* is a taxon of multicellular organisms with a large effective population size, so selective processes are more efficient; mammals – especially primates and *Homo* – are evolutionarily young and have a smaller effective population size, while yeasts are unicellular. If these factors affect general evolutionary properties, such as the selective cost of translation of lowly functional proteins and the probability of fixation by drift, it is possible that they may affect the evolution of *de novo* proteins. In the case of *Homo*, all these factors may be more favorable to the fixation of new *de novo* proteins, and the availability of broad and varied -omics data is unparalleled. It would therefore be an obvious extension to employ a similar approach to investigate possible utORFs and *de novo* proteins in humans.

## Methods

### Comprehensive Drosophila melanogaster ORFeome

To generate the comprehensive database of potential ORFs in *Drosophila melanogaster*, we translated the repeat-masked genome (UCSC Aug. 2014 BDGP Release 6 + ISO1 MT/dm6) and transcriptome (FlyBase, r6.15). We used the repeat-masked genome to control the number of transposable elements and other degenerate candidates. We retained only ORFs of at least 8 residues; for ORFs not containing a canonical start codon, only those of at least 20 residues were retained. We deduplicated potential ORF sequences, leaving 4,583,941 unique potential ORFs.

### Discovery of utORFs in published shotgun MS data

To identify evidence of utORFs, we reanalyzed a published proteome of *D. melanogaster* (PRIDE accession numbers PXD005691 and PXD005713; respectively, whole lifecycle and hourly/bihourly embryogenesis stages; (Casas-Vila et al. 2017)) using MaxQuant v. 1.6.1.0 (Tyanova, Temu, and Cox 2016). To speed computation, we divided analysis of the overall proteome into 4 subsets (**Figure 1 - Source Data 4**). As in (Casas-Vila et al. 2017), we used standard parameters (namely, methionine oxidation and N-terminal acetylation as variable modifications; carbamidomethylation as a fixed modification; tryptic digestion; minimum peptide length 7).

#### Two-round MS search

In the two-round MS search, we used two rounds of analysis per data subset to improve total sensitivity while maintaining an acceptable false discovery rate (**Figure 1A**). In the first “discovery” round, we set the peptide and protein group FDRs in MaxQuant to 0.2, with the protein search sequences set to be the annotated *D. melanogaster* proteome (FlyBase, r6.15), supplemented with the comprehensive potential ORF database. We then identified candidate ORFs from any of the four data subsets (**Figure 1-Source Data 4**) to develop a filtered subset of potential ORFs with possible mass spectrometry evidence. These relaxed parameters allow us to identify many more potential ORFs, but with a corresponding increase in false positive identifications. In the second “verification” round, we set the peptide and protein group FDRs to 0.01, with the protein search sequences set to be the annotated *D. melanogaster* proteome (FlyBase, r6.15), supplemented with only that filtered subset of potential ORFs identified in the first round (around 1000-fold fewer). We subsequently performed additional manual curation of the identified hits to arrive at our final set of utORFs; in brief, we dropped utORFs that were shorter than 14 residues, that had detectable sequence homology to annotated proteins via BLASTP, and the shorter of multiple possible utORFs that were identified from the same protein group by MaxQuant.

#### Evaluation of peptide and protein polymorphisms

Evaluating whether a given peptide could have arisen from a polymorphism is equivalent to calculating the number of substitutions from said peptide to the nearest peptide expected from the annotated proteome. Variations of this problem are well-studied in computer science, and trie data structures are commonly used to solve them efficiently. We used a Python implementation of tries (https://github.com/jfjlaros/dict-trie) to build a trie of all tryptic peptides that would be expected from the annotated *D. melanogaster* proteome. For each peptide uniquely identified from a utORF, we calculated Hamming (the number of substitutions without the possibility of indels) and Levenshtein (the combined number of substitutions and indels) distances.

#### Genomic analysis of utORFs

To map utORF sequences back to their genomic locations, we used BLAT version 36 (Hinrichs et al. 2006) to identify the locations of utORF sequences that were supported in the discovery round. We reformatted the output to a GTF and made certain manual corrections; this file, containing the final locations, is appended as **Figure 1–Source Data 3**. We excluded those loci whose genomic location could not be determined from genomic analyses. For those loci where multiple genomic locations were possible, we used the first location identified by BLAT.

We subsequently classified the genomic locations of these utORF sequences using custom scripts relying on the Bioconductor libraries in R (Huber et al. 2015). Specifically, we classified a location as sense or antisense if any part of the location overlapped with an annotated gene in the appropriate direction. We used the FlyBase r6.15 annotations. We classified a location as intergenic if and only if it was neither sense nor antisense as defined above.

We computed conservation scores by using the 27-way phastCons and phyloP bigWig tracks provided by the UCSC Genome Browser (Hinrichs et al. 2006). For a given genomic location, we calculated the conservation score by averaging the score through the relevant interval.

#### Published sequencing data

We used a number of published sequencing datasets to characterize utORFs. To evaluate developmental stage- and tissue-specific transcription levels, we used modENCODE (modENCODE Consortium et al. 2010) and FlyAtlas2 (Leader et al. 2018) data.

### Gene age inference

#### Multiple sequence alignment and genomic similarity scoring

We used BioPython v. 1.69 (Cock et al. 2009) to subset the UCSC 27-way insect multiple-sequence alignment (MSA) to a smaller MSA that corresponded to the location in the *D. melanogaster* genome of each unannotated translated ORF plus the flanking 9bp on each side. In a pairwise fashion for each species against *D. melanogaster*, we extracted genomic sequences from the MSA, dropped gaps in the sequences, and calculated identity scores.

#### Modified Smith-Waterman protein similarity scoring

We calculated protein similarity scores using the Smith-Waterman algorithm with the BLOSUM62 matrix (Henikoff and Henikoff 1992), no gap or extension penalties, and we normalized by the length of the ORF in *D. melanogaster*. To ensure score stability with respect to minor indels, we calculated protein similarity scores in all three reading frames, keeping only the frame that scored best. In doing so, we considered the entirety of the syntenic block irrespective of the formal boundaries of ORFs in the related species; this allows us to capture the locally-optimal (within the block) similarity and tolerate recent acquisition of nonsense or frameshift mutations.

#### Phylostratigraphy

Using the reference phylogeny corresponding to the UCSC 27-way insect MSA, we inferred approximate gene emergence dates from the pattern of modified Smith-Waterman protein similarity scores. We defined significance as a score greater than 2.5 points. In turn, we estimated the inferred age of a given ORF as the age of the last common ancestor between *D. melanogaster* and the most distant insect species for which a significant modified Smith-Waterman protein similarity score was calculated.

### Latent Class Analysis

We performed latent class analysis (Collins and Lanza 2009) using the *poLCA* R package (Linzer and Lewis 2011). Specifically, we used 100 iterations of a latent class model with no covariates and with factors as follows: inferred gene age; monophyly; genomic location (classified as solely intergenic, antisense, sense, or any combination); phastCons score (binned as [0, 0.2), [0.2, 0.8), and [0.8, 1.0]); length (binned as [0, 20), [20, 50), and [50, infinity); transcription (binned as TPM > 0.1 or ≤ 0.1 from maximum per-sample TPM) tissue specificity (binned as *tau* > 0.8 or ≤ 0.8 from FlyAtlas tissues) developmental specificity (binned as *tau* > 0.8 or ≤ 0.8 from modENCODE developmental stages). We excluded loci with missing values from latent class analysis. We chose the number of classes (five) based on a combination of minimizing the BIC and AIC (**Supplemental File 1D and E**) as well as model interpretation, similar to recommended procedures (Collins and Lanza 2009). For the analysis of all utORFs, models with 6 classes did not converge on a single maximum-likelihood solution, suggesting potential under-identification, and were thus not considered.

We subsequently used the resulting five-class latent class model to predict the posterior class membership probabilities for each classified candidate locus. We used the modal probability to assign class membership. We also performed a parallel analysis of utORFs with canonical start codons, which led to similar conclusions.

### Validation of utORFs

#### Retention time validation

We used AutoRT (Wen et al. 2020) to predict retention times of identified peptides from both annotated FlyBase proteins and utORFs. To train the model, we conducted transfer learning per each source run using the published AutoRT model (originally trained on PXD006109) with 40 epochs on either an NVIDIA K80 GPU or an NVIDIA V100 GPU; the training set consisted of only those annotated FlyBase peptides with posterior error probabilities of 0.01 or smaller as estimated by MaxQuant. We then used the per-run trained models with default parameters to predict the retention times of the peptides resulting from either the annotated FlyBase proteins or utORFs.

#### Empirical validation

We collected protein samples from *D. melanogaster* 0–24-hour embryos (RAL517 strain) for validation of utORFs via mass spectrometry. In brief, we homogenized dechorionated embryos with a mortar and pestle in ice-cold lysis buffer (50mM Tris HCl, pH7.5; 137 mM NaCl; 0.25% NP-40; Roche cOmplete protease inhibitors) before centrifugation at 18,000 rpm at 4°C and filtration through a 0.45 micron PVDF membrane.

Purified protein was acetone-precipitated, reduced, and alkylated before digestion with LysC and trypsin, acidified, and solid-phase extraction. The resulting peptides were analyzed by LC-MS/MS (120min gradient, 25cm EasySprayer column in high res./high mass accuracy mode) on a ThermoFisher Lumos instrument. In addition, the embryo sample was analyzed on the same instrument by PRM-MS, targeting a set of 16 tryptic peptides. We manually chose these targeted peptides based on suitability of the peptide for targeting (i.e., few/no missed cleavages or modifications), expression of the utORF in embryos, and the strength of the prior detection from the Casas-Vila datasets.

We analyzed the shotgun data using ProteomeDiscoverer 1.4 and searched via Mascot against the Uniprot D. melanogaster database concatenated with common contaminants and our set of utORFs with a 1% peptide FDR. We analyzed the targeted data using SkylineMS (v. 21.2.0.369 (2efacf038)). We deposited the raw data in PRIDE under accession PXD032197.

We additionally reanalyzed published ribosome profiling data downloaded from the NCBI Sequence Read Archive (SRA) project PRJNA306373 (Zhang et al. 2018) and project SRP254283 (Patraquim et al. 2020). For the Zhang dataset, we downloaded raw reads from the above SRA accessions, aligned them to the genome using hisat2 v.2.2.1 (Kim et al. 2019), and calculated TPM using stringtie v.1.3.4d (Pertea et al. 2016). We handled the Patraquim dataset similarly, except that we additionally trimmed the raw reads using trimmomatic 0.38 (Bolger, Lohse, and Usadel 2014) and filtered out reads aligning to annotated RNAs using bowtie2 v. 2.3.5 (Langmead and Salzberg 2012) before alignment. For each utORF, we calculated the maximum TPM observed across either dataset and called the utORF as supported if the TPM was greater than 0.2.

To identify utORFs with statistically significant framing of ribosome-profiling reads, we used a binomial test as in Patraquim et al. (2020). We aligned the trimmed, filtered reads to the set of sequences corresponding to our utORFs, plus an additional 30bp up- and down-stream, using bowtie v. 1.3.1 (Langmead et al., 2009) with options --tryhard --suppress 1,6,7,8. We analyzed the resulting aligned footprints using the R package riboSeqR v 1.20.0. For each utORF with at least two unique footprints, we calculated the binomial probability of observing the number of footprints in the utORF’s frame versus the other two frames at an uncorrected alpha of 0.05. We pooled footprints across replicates and across read lengths for a given utORF in each sample.

## Data Availability

Raw MS data are deposited in PRIDE under accession number PXD032197. Relevant scripts and intermediate files can be found in our Github repository https://github.com/LiZhaoLab/utORF_mass_spec.

## Acknowledgements

We thank members of the Zhao laboratory for helpful discussions during the work. We thank Henrik Molina and the Proteomics Resource Center at the Rockefeller University for the help with mass spectrometry data.

## Supplemental Figure Legends

**Figure 3–figure supplement 1:**
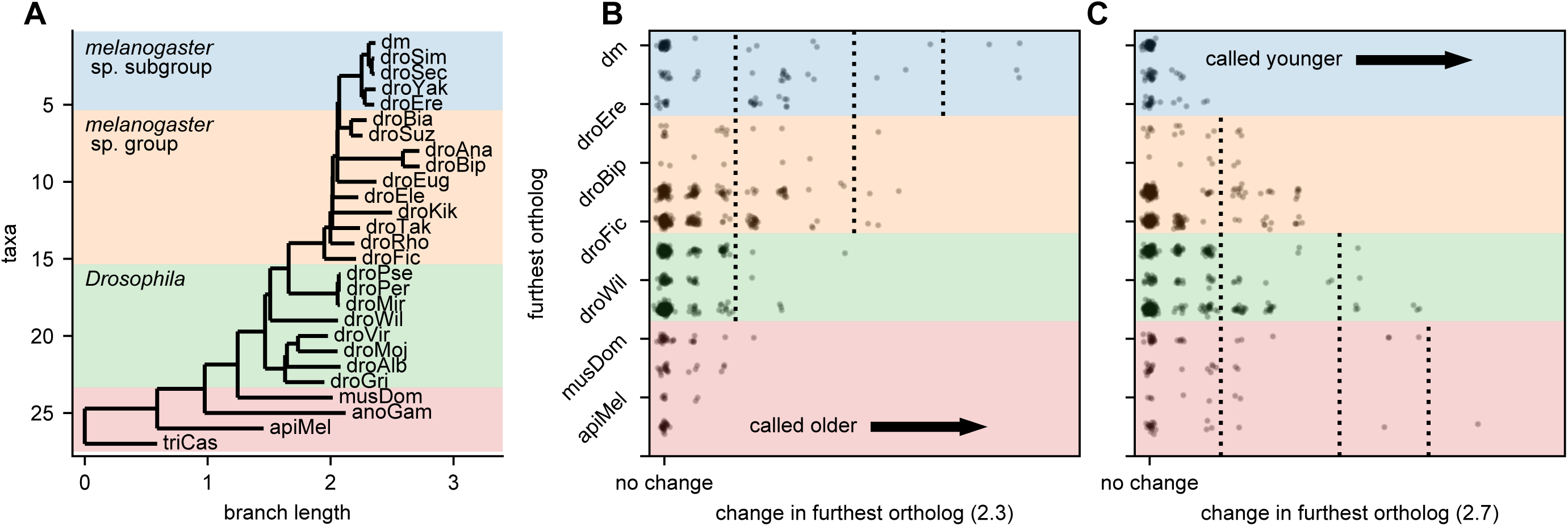
Robustness of gene age inferences with respect to significance threshold. **A**: Figure 3A, but also showing the *melanogaster* species subgroup, species group, and *Drosophila* taxa. **B**: Change in furthest significant ortholog (using a significance threshold of 2.3 instead of 2.5). Coloring corresponds to panel A. Lowering the threshold causes more orthologs of utORFs to be called as significant, increasing the inferred gene age. **C**: As in panel B, but with a significance threshold of 2.7 instead of 2.5. Increasing the threshold causes fewer orthologs of utORFs to be called as significant, reducing the inferred gene age.

**Figure 4–figure supplement 1:**
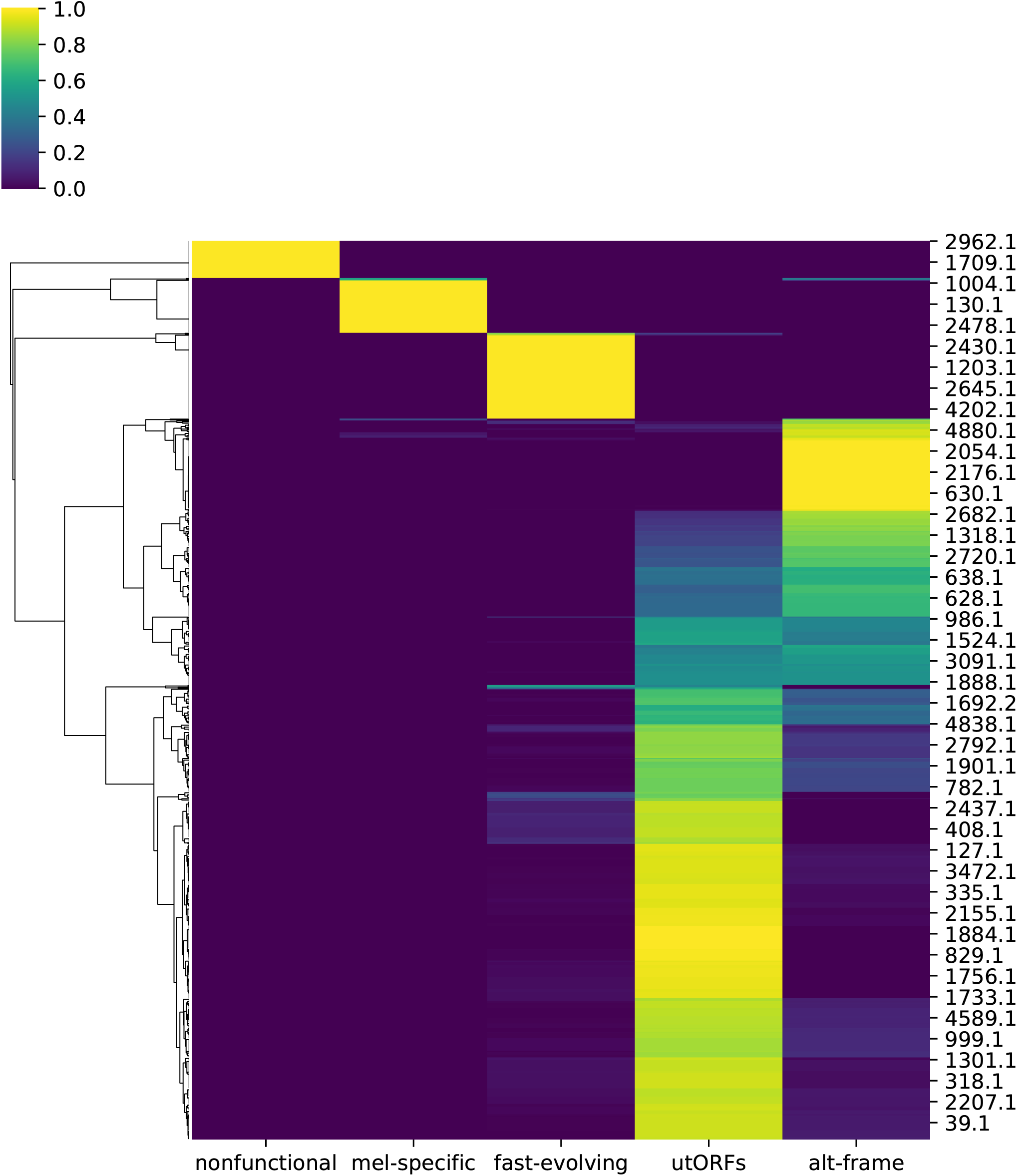
Posterior probabilities per unannotated translated ORF of class membership inferred from latent class analysis for all utORFs.

**Figure 5–figure supplement 1:**
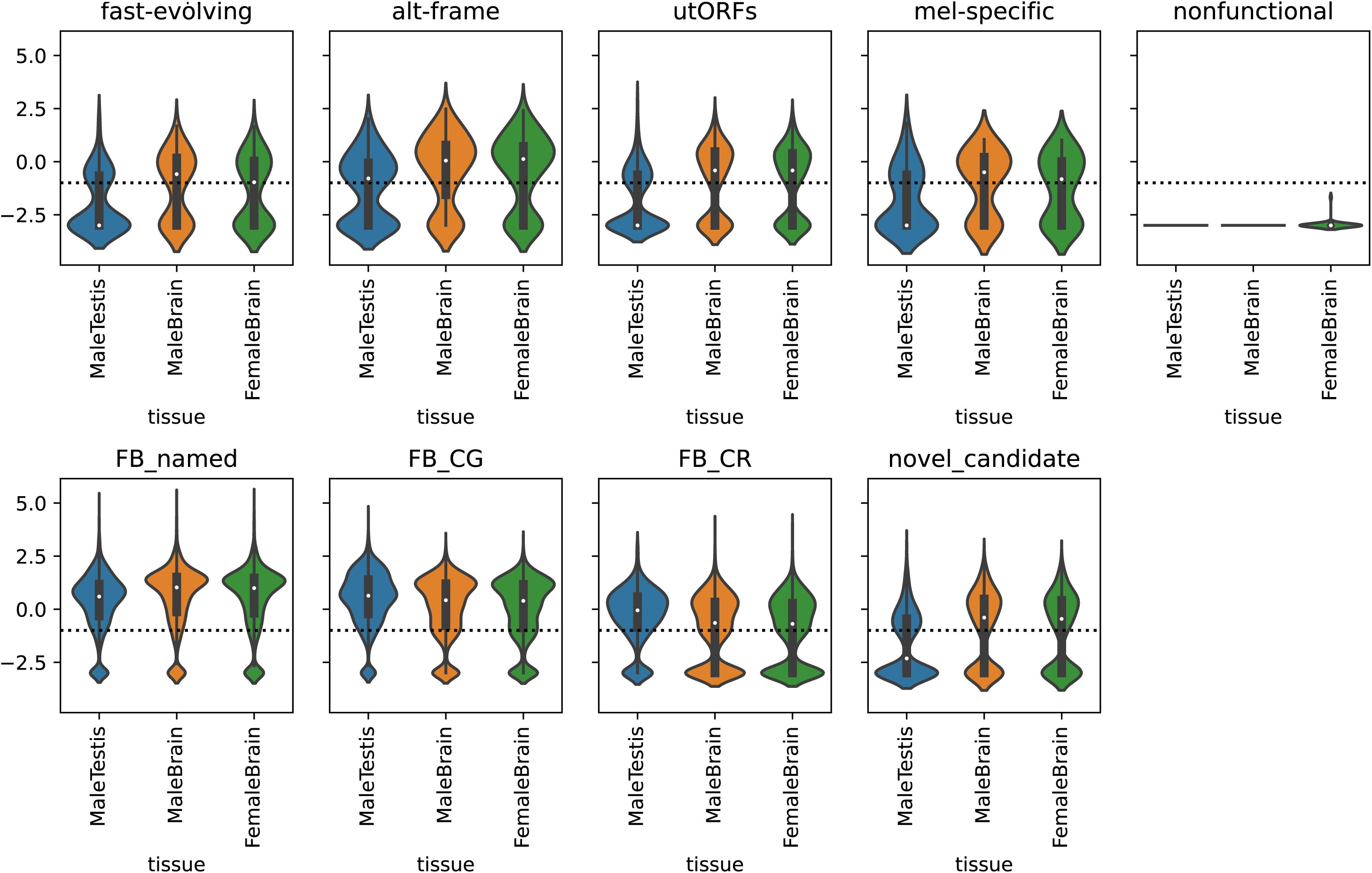
Transcription of utORFs in selected tissues. Top panel: utORFs, separated by inferred LCA class. Mean TPMs across the given tissue in FlyAtlas2 are log10-transformed with a pseudocount of 1E-3. Horizontal line marks an expression threshold of TPM 0.1. Bottom panel: As top, but for, in order, annotated FlyBase genes, separated between those with names, those with “CG” identifiers, and those with “CR” identifiers, and all utORFs.

**Figure 4–figure supplement 2:**
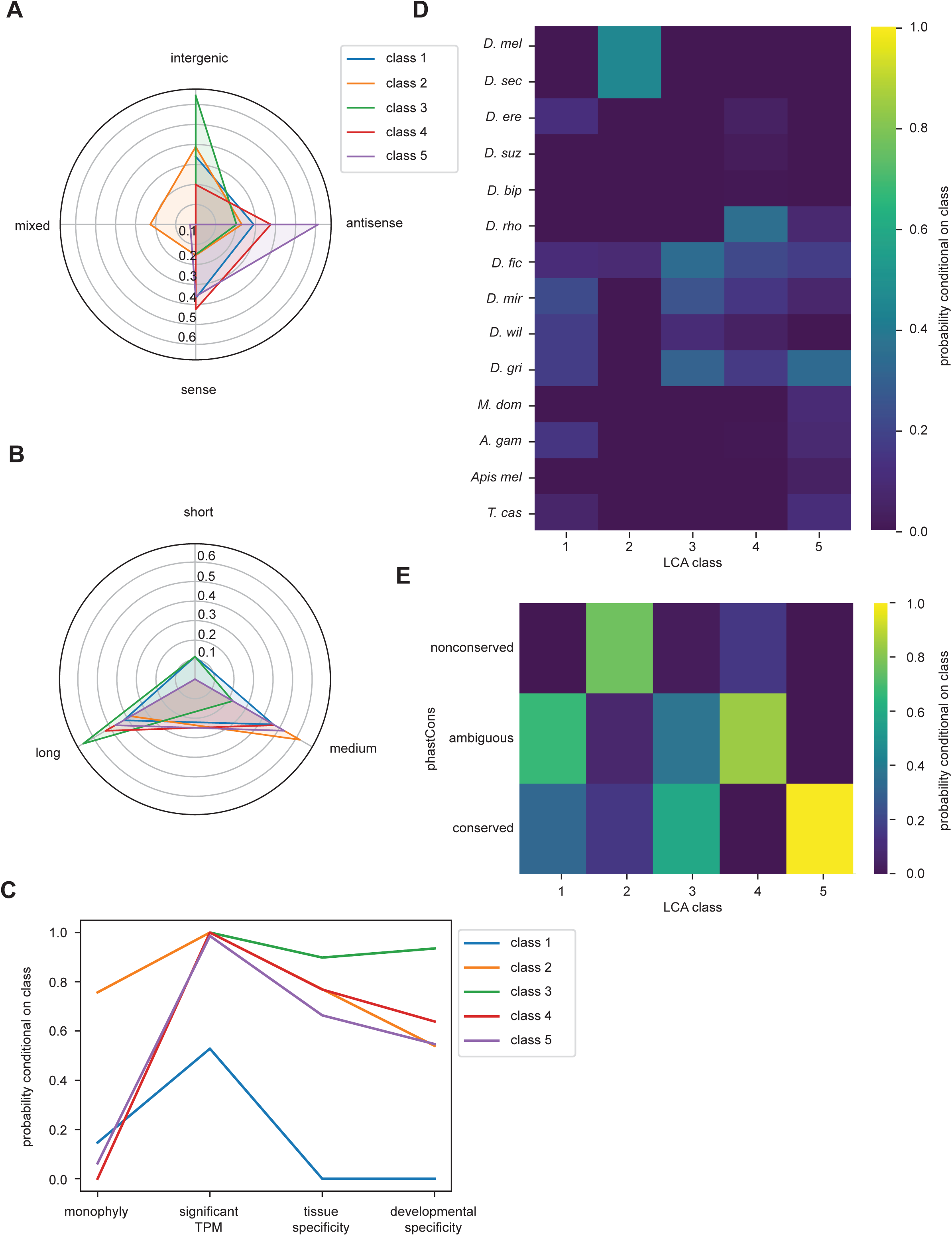
Latent class analysis of utORFs with canonical start sites reveals differences between classes. **A-E:** same as Figure 4, but examining utORFs with canonical start sites.

**Figure 4–figure supplement 3:**
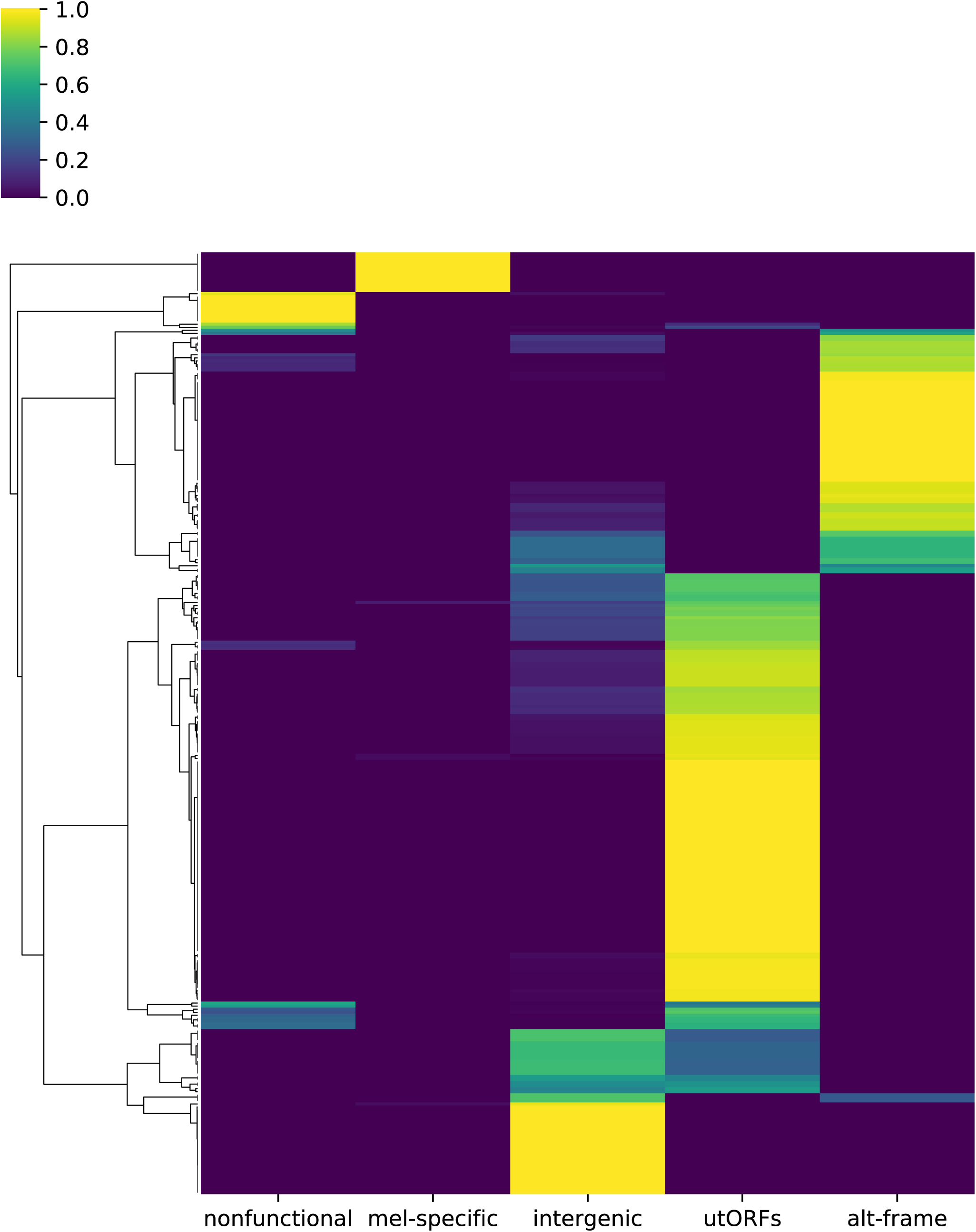
Posterior probabilities per unannotated translated ORF of class membership inferred from latent class analysis for utORFs with canonical start sites.

**Figure 5–figure supplement 2:**
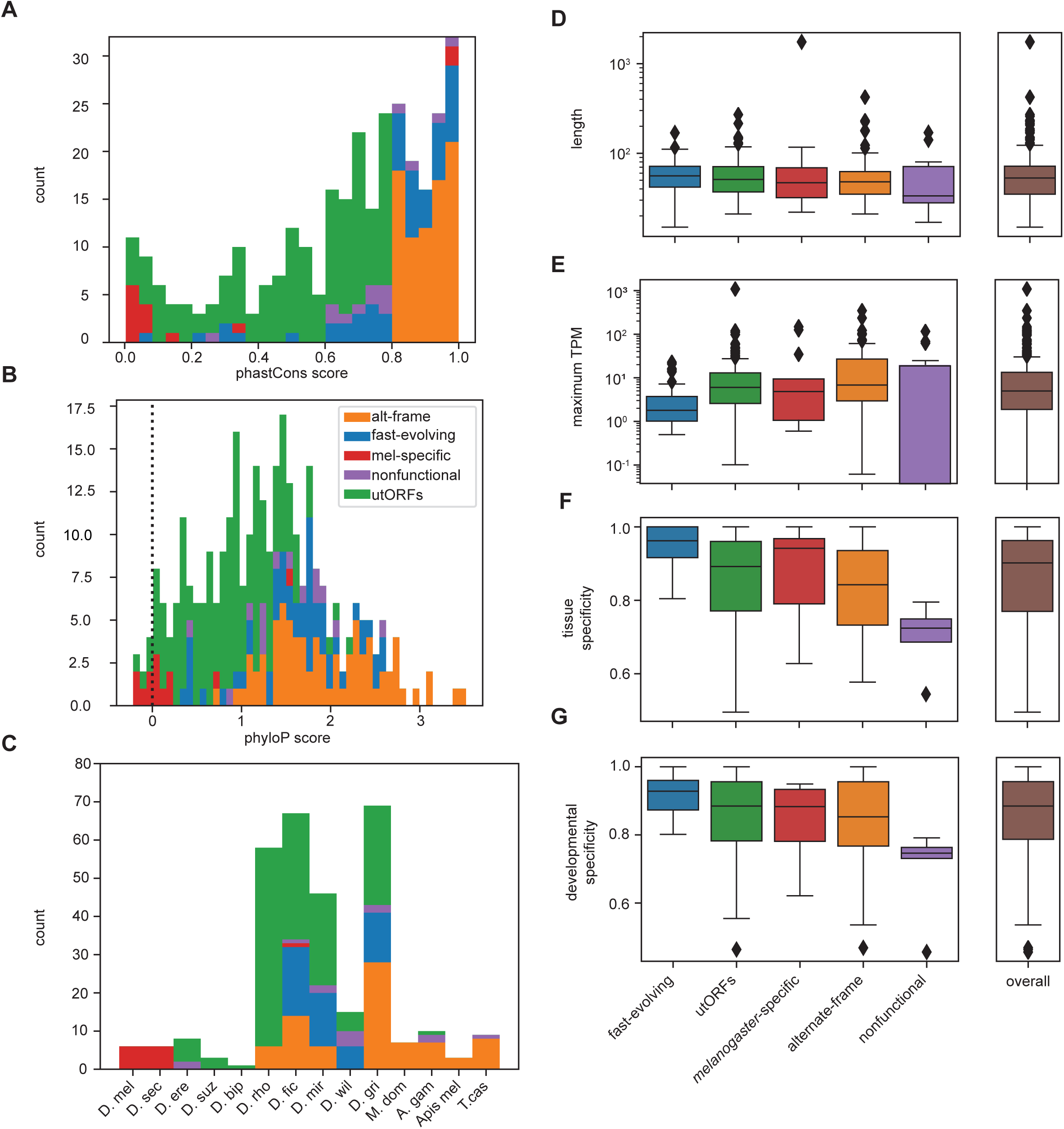
Differences between inferred classes for utORFs with canonical start sites recapitulate expected trends. **A-G**: same as Figure 5, but examining utORFs with canonical start sites.

**Figure 7–figure supplement 1:**
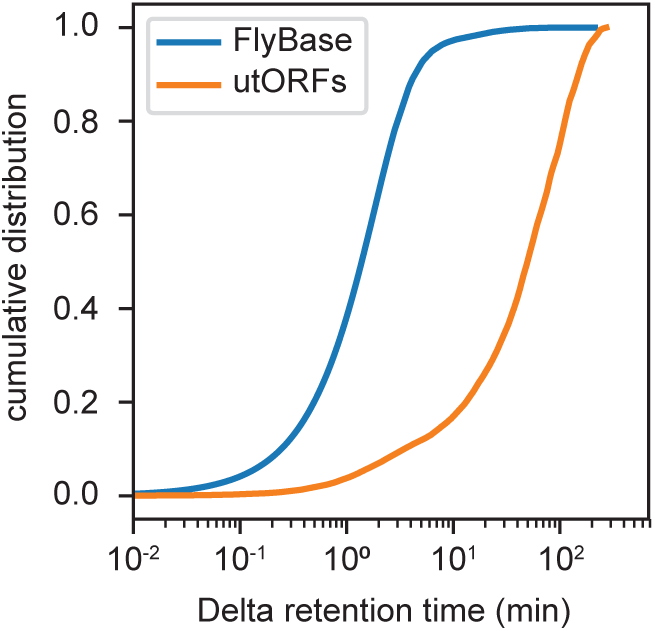
Cumulative distribution of differences between observed and predicted retention times for every peptide-spectrum match of peptides supporting all annotated FlyBase proteins (blue) and all utORFs (orange).

**Figure 7–figure supplement 2:**
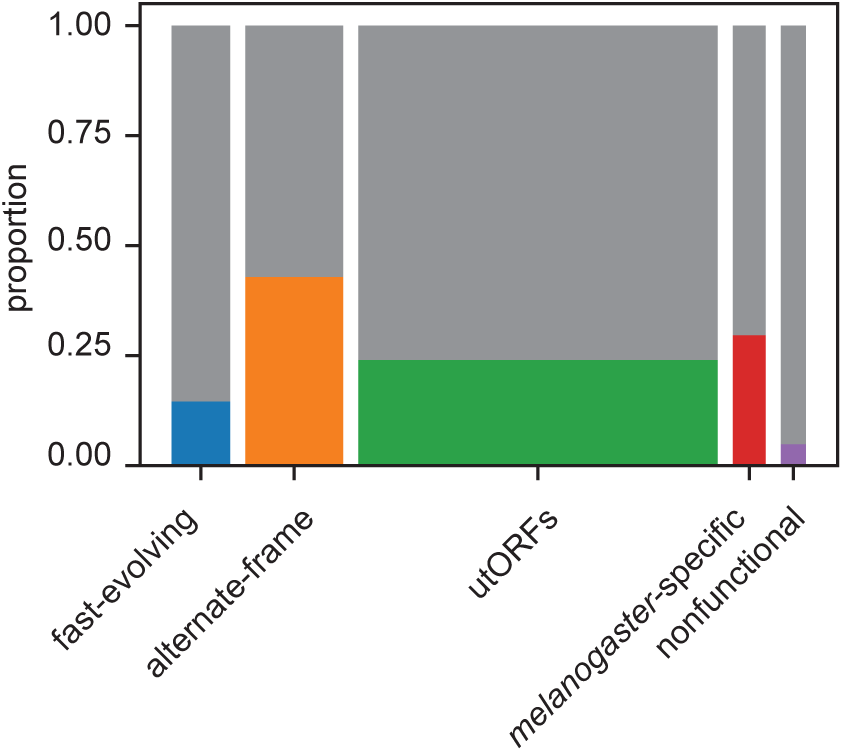
Proportion of utORFs by inferred class with supporting evidence from ribosome profiling. Box widths correlate with size of class (Table 1).

**Supplemental File 1A:** Two-round analysis improves identifications.

**Supplemental File 1B**: Latent class analysis of unannotated translated ORFs with canonical start sites.

**Supplemental File 1C**: AutoRT-supported identifications by inferred gene class.

**Supplemental File 1D**: Summary statistics used for selecting the number of latent classes when considering all utORFs. Five latent classes were chosen as a balance of minimizing the AIC and BIC and model interpretation. AIC: Akaike information criterion; BIC: Bayesian information criterion; G^2^: likelihood-ratio statistic; Chi-sq: Chi-squared. * : Models with 6 classes did not converge on a single maximum-likelihood solution within 100 replicates.

**Supplemental File 1E**: Summary statistics used for selecting the number of latent classes when considering only utORFs with canonical start codons. Five latent classes were chosen as a balance of minimizing the AIC and BIC and model interpretation. AIC: Akaike information criterion; BIC: Bayesian information criterion; G^2^: likelihood-ratio statistic; Chi-sq: Chi-squared.

## Source Data Files

Figure 1–Source Data 1 (csv): all utORFs sequences.

Figure 1–Source Data 2 (csv): utORF supporting peptides.

Figure 1–Source Data 3 (gtf): utORF locations.

Figure 4 –Source Data 1 (csv): utORF inferred LCA classes.

Figure 7–Source Data 1 (xlsx): MASCOT search results from embryo MS data.

Figure 7–Source Data 2 (csv): rank of potentially biologically significant targets.

Figure 1–Source Data 4 (xlsx) - dataset subset mappings.

