## Supplemental File 1 for "Protein evidence of unannotated ORFs in *Drosophila* reveals unappreciated diversity in the evolution of young proteins"

Systematic identification of unannotated ORFs in Drosophila reveals evolutionary heterogeneity

**Supplemental Table 1A:** Two-round analysis improves identifications.

| Methodology | Total Proteins | Contaminants | Decoy IDs | Annotated IDs | Unannotated IDs |
| --- | --- | --- | --- | --- | --- |
| Standard single-round, FDR 0.01 | 11578 | 65 | 440 | 10936 | 137 |
| Two-step “discovery” round, FDR 0.2 | 28483 | 49 | 5220 | 18182 | 5032 |
| Two-step “verification” round, FDR 0.01 | 14084 | 45 | 252 | 11666 | 2121 |

**Total Proteins**: total protein groups identified; **Contaminants**: identifications from contaminant species (e.g. yeast, bacteria, etc.); **Decoy IDs**: identifications of reverse sequence decoys (Cox and Mann, 2008); **Annotated IDs**: groups containing an annotated protein (FlyBase r6.15); **Unannotated IDs**: groups solely containing unannotated candidates; later filtering as in **Methods** (i.e. BLAST searches for duplicates, etc.) reduced these from 2121 to 993 candidates. Note that even at the end of the verification round, with the same FDR threshold and a comparable number of decoy matches, the number of unannotated matches is substantially greater.

**Supplemental Table 1B**: Latent class analysis of unannotated translated ORFs with canonical start sites.

| Class | Interpretation | Estimated Percent | Number |
| --- | --- | --- | --- |
| 1 | putatively nonfunctional loci | 4.55% | 14 |
| 2 | melanogaster-specific ORFs | 4.22% | 13 |
| 3 | intergenic ORFs | 16.56% | 51 |
| 4 | unannotated ORFs | 49.03% | 151 |
| 5 | alternative frame ORFs | 25.65% | 79 |

**Supplemental Table 1C**: AutoRT-supported identifications by inferred gene class.

| class | supported | not supported | proportion |
| --- | --- | --- | --- |
| unannotated ORF | 158 | 433 | 26.7% |
| alternative-frame | 48 | 113 | 29.8% |
| fast-evolving | 25 | 71 | 26.0% |
| melanogaster-specific | 17 | 37 | 31.5% |
| Nonfunctional loci | 6 | 35 | 14.6% |

**Supplemental Table 1D**: Summary statistics used for selecting the number of latent classes when considering all utORFs.

| Number of Classes | AIC | BIC | G^2^ | Chi-sq |
| --- | --- | --- | --- | --- |
| 2 | 13202.7 | 13442.44 | 1806.049 | 11173.71 |
| 3 | 13041.25 | 13403.31 | 1594.601 | 7488.03 |
| 4 | 12931.12 | 13415.5 | 1434.47 | 4956.729 |
| 5 | 12857.13 | 13463.81 | 1310.472 | 4788.56 |
| 6* | 12820.76 | 13549.76 | 1224.104 | 4155.138 |

Five latent classes were chosen as a balance of minimizing the AIC and BIC and model interpretation. AIC: Akaike information criterion; BIC: Bayesian information criterion; G^2^: likelihood-ratio statistic; Chi-sq: Chi-squared *: Models with 6 classes did not converge on a single maximum-likelihood solution within 100 replicates.

**Supplemental Table 1E**: Summary statistics used for selecting the number of latent classes when considering only utORFs with canonical start codons.

| Number of Classes | AIC | BIC | G^2^ | Chi-sq |
| --- | --- | --- | --- | --- |
| 2 | 4020.45 | 4203.225 | 797.163 | 12055.97 |
| 3 | 3983.328 | 4259.356 | 710.041 | 4596.399 |
| 4 | 3978.968 | 4348.247 | 655.680 | 3333.585 |
| 5 | 3975.269 | 4437.801 | 601.982 | 1720.968 |
| 6 | 3978.989 | 4534.774 | 555.702 | 1498.315 |

Five latent classes were chosen as a balance of minimizing the AIC and BIC and model interpretation. AIC: Akaike information criterion; BIC: Bayesian information criterion; G^2^: likelihood-ratio statistic; Chi-sq: Chi-squared.
